# An O-GlcNAc transferase pathogenic variant that affects pluripotent stem cell self-renewal

**DOI:** 10.1101/2023.03.13.531514

**Authors:** Michaela Omelková, Christina Dühring Fenger, Marta Murray, Trine Bjørg Hammer, Veronica M. Pravata, Sergio Galan Bartual, Ignacy Czajewski, Allan Bayat, Andrew T. Ferenbach, Marios P. Stavridis, Daan M. F. van Aalten

## Abstract

O-linked β-N-acetylglucosamine (O-GlcNAc) transferase (OGT) is an essential enzyme that modifies proteins with O-GlcNAc. Inborn *OGT* genetic variants were recently shown to mediate a novel type of Congenital Disorder of Glycosylation (OGT-CDG) which is characterized by X-linked intellectual disability (XLID) and developmental delay. Here, we report an OGT^C921Y^ variant which co-segregates with XLID and epileptic seizures, and results in loss of catalytic activity. Colonies formed by mouse embryonic stem cells carrying OGT^C921Y^ show decreased levels of protein O-GlcNAcylation accompanied by decreased levels of Oct4, Sox2 and extracellular alkaline phosphatase (ALP), implying reduced self-renewal capacity. These data establish a link between OGT-CDG and embryonic stem cell self-renewal, providing a foundation for examining the developmental aetiology of this syndrome.

**Summary statement:** We show that the C921Y O-GlcNAc transferase variant found in patients with intellectual disability leads to a defect in pluripotent stem cell self-renewal and decreased levels of stem cell markers.

## Introduction

Intellectual disability (ID) is a cognitive handicap that affects up to 1 % of the general population (Maulik et al., 2011; McKenzie et al., 2016). ID is characterized by reduced cognitive function (IQ < 70) and adaptive behaviour diagnosed before the 18^th^ year of life (Tassé et al., 2016). It is estimated that 40 % of ID cases can be attributed to genetic causes (Posada De La Paz et al., 2017). To date, > 2000 genes have been implicated in ID aetiology (Firth and Wright, 2011; PanelApp Genomics England, 2020) and 141 of these are located on the X chromosome (Tejada and Ibarluzea, 2020), which encodes a disproportionate number of genes involved in cognitive processes (Lubs et al., 2012; Skuse, 2005; Zechner et al., 2001). Indeed, due to hemizygosity for X-linked genes, more males than females suffer from ID (Posada De La Paz et al., 2017). Based on aetiology and clinical presentation, ID comprises a heterogenous group of neurodevelopmental conditions that can either be syndromic (i.e., a part of a clinically defined set of symptoms forming a syndrome) or non-syndromic.

Congenital Disorders of Glycosylation (CDG) is an umbrella term for inborn defects in glycosylation enzymes that lead to ID in 60 – 70 % of patients (Wolfe and Krasnewich, 2013). Recently, a novel type of CDG (OGT-CDG) was diagnosed in patients carrying pathogenic variants in O-linked β-*N*-acetylglucosamine (<u>O</u>-<u>G</u>lcNAc) transferase (*OGT*) (Pravata et al., 2020b). To date, 17 pathogenic OGT-CDG variants have been identified through whole genome sequencing, and several of these have been characterized clinically and biochemically (Pravata et al., 2019; Pravata et al., 2020a; Selvan et al., 2018; Vaidyanathan et al., 2017; Willems et al., 2017). In addition to ID and delayed achievement of developmental milestones, patients with OGT-CDG commonly present with musculoskeletal problems, brain malformations, eye abnormalities, dysmorphic features and language problems (Pravata et al., 2020b). Furthermore, heart anomalies, immune system defects, genital abnormalities and digestive problems have also been reported in individual patients. OGT-CDG thus appears to affect the development and function of multiple organ systems. The mechanisms through which OGT variants mediate ID remain to be established, although several hypotheses have been put forward (Pravata et al., 2020b).

OGT is a 117 kDa enzyme encoded by the *OGT* gene located on the X chromosome and expressed ubiquitously in human tissues (Kreppel et al., 1997; Lubas et al., 1997). OGT catalyses O-GlcNAcylation, an essential, dynamic modification of nucleocytoplasmic proteins with a single *O*-linked GlcNAc sugar moiety on serine and threonine residues (Kreppel et al., 1997; Torres and Hart, 1984). As opposed to other post-translational modifications that are generally mediated by protein families, O-GlcNAcylation is controlled by only two enzymes with opposing function, OGT and <u>O</u>-<u>G</u>lcNAc hydrol<u>a</u>se (OGA), that attach and remove the GlcNAc moiety, respectively (Gao et al., 2001; Wells et al., 2002). The sugar nucleotide donor for O-GlcNAcylation, UDP-GlcNAc, is synthesized from glucose by the hexosamine biosynthetic pathway and thus O-GlcNAcylation is thought to serve as a nutrient sensor (Hart, 2015; Medford et al., 2013). Since 1984, when O-GlcNAcylation was first discovered in lymphocytes (Torres and Hart, 1984), modification of O-GlcNAc proteins has been shown to be ubiquitous across all human tissues (Wulff-Fuentes et al., 2021). O-GlcNAcylation has been implicated in a range of biological mechanisms such as cellular metabolism (Slawson et al., 2010), gene expression and regulation (Krause et al., 2018; Sakabe et al., 2010; Streubel et al., 2017; Tan et al., 2021), cell division (Drougat et al., 2012; Lefebvre et al., 2004; Leturcq and Lefebvre, 2017; Sakabe and Hart, 2010) and stem cell differentiation (Elena-Herrmann et al., 2020; Hao et al., 2019; Zhang et al., 2019). Moreover, O-GlcNAcylation is considered essential for survival as embryonic ablation of *Ogt* in mice (Shafi et al., 2000) and zebrafish (Webster et al., 2009) is lethal, while loss of Oga activity leads to perinatal death in mice (Keembiyehetty et al., 2015; Muha et al., 2021; Yang et al., 2012).

OGT is composed of 13.5 N-terminal tetratricopeptide repeats (TPRs) that mediate substrate recognition and protein - protein interactions (Iyer and Hart, 2003; Jínek et al., 2004; Rafie et al., 2017) and a globular catalytic domain that catalyses O-GlcNAcylation. In addition to acting as an essential glycosylation enzyme, OGT also proteolytically activates transcriptional co-regulator <u>H</u>ost <u>C</u>ell <u>F</u>actor 1 (HCF1) (Capotosti et al., 2011). HCF1 is heavily glycosylated and cleaved by OGT in the same active site, however, glycosylation and proteolysis occur through separate mechanisms (Kapuria et al., 2018; Lazarus et al., 2013). Data suggest that HCF1 may bind a large proportion of promoters in the human genome, acting as an important cell cycle progression and mitochondrial biogenesis regulator (Michaud et al., 2013). Interestingly, *HCFC1* is itself a well-known XLID gene, mutations of which are thought to alter neural stem cell maintenance and differentiation (Castro et al., 2020; Huang et al., 2012; Jolly et al., 2015).

There is growing evidence to suggest that O-GlcNAcylation is essential for embryonic stem cell (ESC) maintenance (Shafi et al., 2000) and embryogenesis (Jang et al., 2012; Shafi et al., 2000; Webster et al., 2009; Yang et al., 2012; Zhang et al., 2019; Zhu et al., 2020). Cellular O-GlcNAcylation decreases during neuronal differentiation (Liu et al., 2012), suggesting a link between dynamic O-GlcNAc cycling and tissue development. Furthermore, OGT O-GlcNAcylates transcription factors that play an important role in pluripotency and stem cell function, including <u>Oc</u>tamer binding transcription factor <u>4</u> (Oct4) (Constable et al., 2017), <u>S</u>RY (sex determining region Y)-b<u>ox</u> <u>2</u> (Sox2) and <u>S</u>ignal transducer and <u>a</u>ctivator of transcription <u>3</u> (STAT3) (Li et al., 2017). STAT3 is a member of the JAK/STAT signalling pathway that is activated by pluripotency stimulus <u>L</u>eukemia Inhibiting <u>F</u>actor (LIF) in mouse ESCs (mESCs). Here, we report three brothers affected by OGT-CDG that carry a novel variant, OGT^C921Y^, in the catalytic core of OGT that is absent in their healthy brother. The variant has not been previously reported, neither in the healthy population (gnomAD database) nor in patients (HGMD database). We show that the OGT^C921Y^ variant possesses decreased glycosyltransferase activity *in vitro* and in an mESC model. Strikingly, a knock-in of OGT^C921Y^ results in abrogated self-renewal in mESCs as shown by decreased expression of alkaline phosphatase (ALP), Oct4 and Sox2 during a clonogenic assay. These results suggest that the role of OGT in ESC self-renewal and pluripotency may contribute to clinical signs seen in OGT-CDG patients.

## Results and Discussion

### Three brothers with intellectual disability carry an inherited catalytic OGT variant absent in their healthy brother

Four male siblings were born to a healthy non-consanguineous couple shown as family members II.5 and II.6 on the family pedigree (Fig. 1A). The first child was born in 1960 and is healthy. The second child, currently a 58-year-old male shown as proband III:2 on the family pedigree, was born at 38 weeks of gestation after a normal pregnancy. Birth weight was 2950 gram (2^nd^ - 9^th^ centile) and birth length was 52 cm (50^th^ - 75^th^ centile). Apgar scores at birth are unknown. There were no concerns following birth and the patient was discharged. Following the first two years of life, he showed delay in reaching developmental milestones, especially in areas of speech and language development. He learned his first words at the age of 14 months, and he is now able to speak in short sentences with five to seven words and understand simple instructions. At the age of twelve months, he could walk and reach out with palmar grasp, transfer objects, and put them into his mouth. Later in life, the patient presented with autistic features. He attended a special needs school, and he currently lives in a sheltered home. He underwent an operation for an inguinal hernia at the age of 48 years. Brain MRI was never performed. He has experienced at least two generalised tonic clonic seizures. The first seizure occurred around the age of 40 years. At seizure onset, his EEG showed background slowing, but without any clear interictal epileptiform abnormalities. After the second seizure, a daily treatment with oxcarbazepine was prescribed and he has since had a good seizure control. Dysmorphic features include an oval face, narrow, long and pear-shaped nose with a high nasal bridge, thin upper lip, high arched palate, large ears, sparse eyebrows, thin and short fingers with distal squaring (Fig. 1B, Fig. S1). At the age of 58, the patient has osteoporosis and scoliosis. He has been diagnosed with both a short stature (166 cm, - 1.48 SD) and a head circumference of 55.3 cm (+ 0.12 SD). Cranial nerve examination was normal. Limb examination showed neither rigidity nor tremors. The power in the limbs was five out of five (MRC Scale for Muscle Strength) and the deep tendon reflexes were normal. He presented with a slow, shuffling gait.

**Figure 1:**
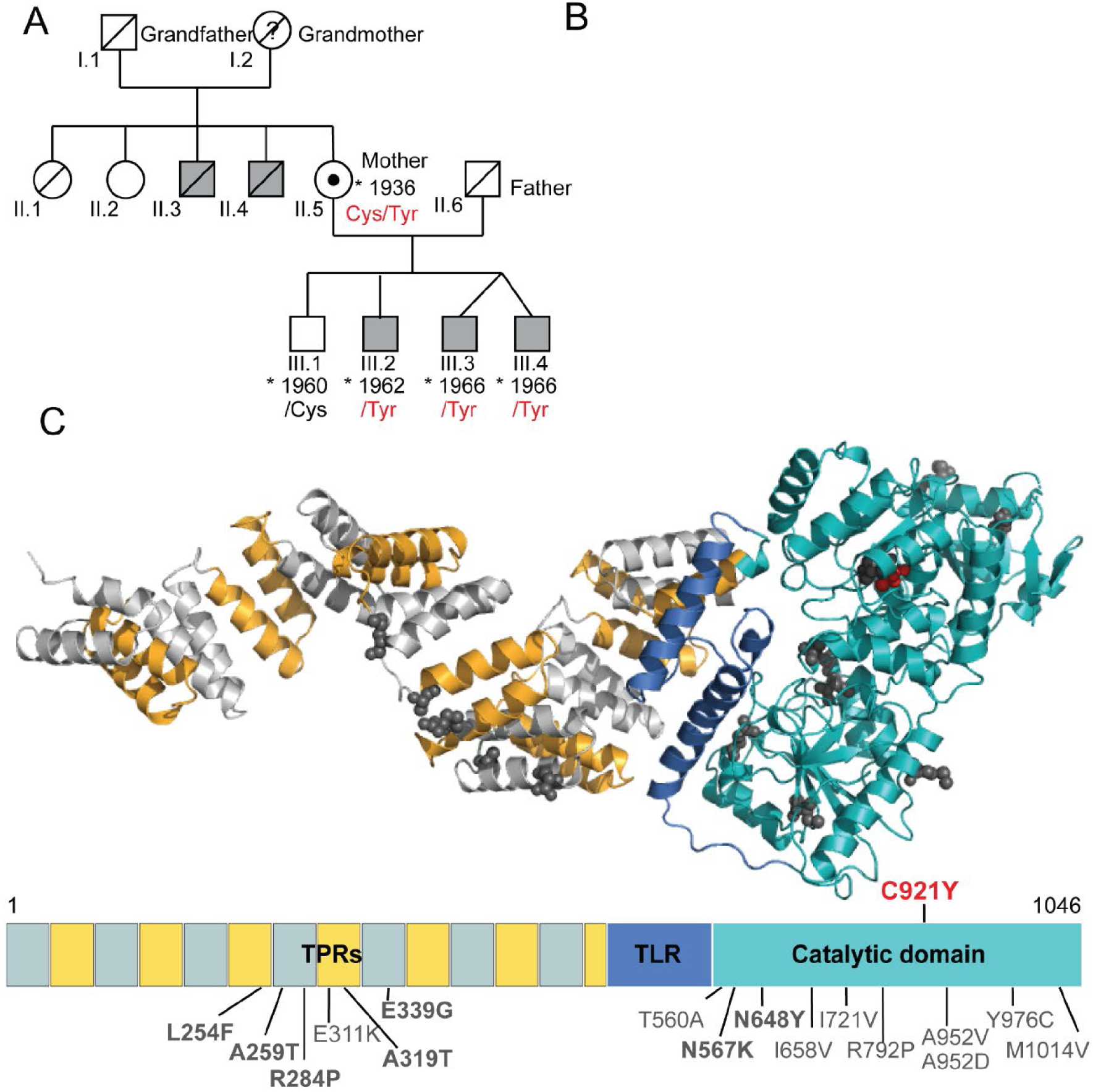
Brothers with intellectual disability carry an inherited variant in the catalytic core of OGT that is absent in their healthy brother. **(a)** Pedigree of the affected patients carrying the OGT^C921Y^ variant. Grey shapes denote individuals affected by intellectual disability. Where available, the OGT variant at position 921 is provided. Black dot indicates a female carrier of the OGT^C921Y^ variant. **(b) *Figure panel censored for BioRxiv submission.*** Facial photographs of two brothers carrying OGT^C921Y^ variant with corresponding family tree annotation. Patient III.2 is depicted at age of 58 and patients III.3 and III.4 are depicted at age of 54. **(c)** 3D model of full-length human OGT and the corresponding linear domain structure. TPRs are depicted in alternating gray and yellow, the TLR in dark blue and the catalytic core in cyan. Previously known OGT variants leading to ID and their corresponding residues are highlighted in dark gray. OGT^C921Y^ is highlighted in red. UDP and acceptor peptide molecules are shown yellow and magenta sticks respectively. The 3D model was produced from OGT catalytic core Protein Data Bank code 5C1D, TPR domain Protein Data Bank code 1W3B.

The mother later gave birth to monozygotic male twins, here referred to as twin 1 and twin 2 (III:3 and III:4 in pedigree) at 38 weeks of gestation following a normal pregnancy. Conception was unassisted. Twin 1 (III:3) weighed 2150 gram at birth (< 0.4^th^ centile) and had a birth length of 48cm (2^nd^ - 9^th^ centile), while twin 2 (III:4) weighed 1650 gram (< 0.4^th^ centile) and measured 46 cm (0.4^th^ – 2^nd^ centile). Apgar scores are unavailable. While twin 1 was breastfed and discharged within a week following the birth, twin 2 was tube fed for two weeks. Similar to proband III:2, the twins also showed delay in reaching developmental milestones, especially in areas of speech and language development. Twin 2 never learned to speak and twin 1 is able to pronounce a few words. The twins have autism and communicate using simple signs and by pointing. They are able to understand very simple instructions. Both make high pitched sounds, have repetitive mannerisms and clap their hands together if excited. In addition, twin 2 has self-injurious behaviour. Their motor development was also affected; although both twins learned to walk by the age of 18 months, they were diagnosed with significant gross motor difficulties and clumsiness. They have attended a special needs nursery and school and they currently live in a sheltered home. Neither of the twins underwent a brain MRI. While twin 1 has not had an epileptic seizure, twin 2 has experienced his first generalized tonic clonic seizure at around 40 years of age. Over the subsequent eleven years, he had a total of four identical epileptic seizures.

At the time of the first seizure, the EEG in twin 2 showed background slowing, but without any clear interictal epileptiform abnormalities. After the second seizure, a daily treatment with carbamazepine was prescribed. Data on dysmorphic features from childhood were not available, but both twins currently present with an oval face, up-slanting palpebral fissures, narrow, long and pear-shaped nose with a high nasal bridge, thin upper lip, hypotrichosis and thin fingers (Fig. 1B, Fig. S1). Twin 2 has also been diagnosed with scoliosis, short stature (165 cm, - 2.1 SD) and osteoporosis. In addition, twin 2 is without any rigidity or tremors of the extremities but has a slow shuffling gait. Limited clinical data is available for twin 1.

The three brothers III:2, III:3 and III:4 thus appeared to suffer from an inherited developmental defect which was not present in their eldest healthy brother (III:1). Family history revealed that two maternal uncles (II.3 and II.4) also suffered from intellectual disability and were wheelchair bound at older age, suggesting an X-linked recessive inheritance pattern. Whole genome sequencing of patient III:2 and subsequent Sanger analysis of the *OGT* gene in samples from all three remaining brothers and their mother were performed. These analyses revealed that the affected siblings III:2, III:3 and III:4 carried a guanine to adenine substitution at 2762^nd^ nucleotide (c.2762G>A) in the *OGT* gene that translates as cysteine to tyrosine substitution at position 921 in OGT (c.2762G>A p.(C921Y), NM_181672.2, OGT^921Tyr^) inherited from their heterozygous mother (OGT^921Cys/Tyr^). This OGT variant has not been previously reported and it is classified as likely pathogenic according to the ACMG criteria (criteria PM1 PM2 PP1 PP2 PP3 PP4) (Richards et al., 2015). The OGT^C921Y^ variant was absent in the healthy brother (II:1, OGT^921Cys^), suggesting that this single point mutation in OGT causes intellectual disability in patients II:2, II:3 and II:4. Thus, we have identified three brothers with intellectual disability who carry an inherited catalytic OGT variant absent in healthy brother. The clinical observations are consistent with previous clinical descriptions of OGT-CDG patients and add considerable knowledge to the currently limited information on adult OGT-CDG patients (Pravata et al., 2020b). For example, despite an established association between decreased O-GlcNAcylation and Alzheimer’s disease (Liu et al., 2004; Liu et al., 2009; Park et al., 2021), these patients do not present with signs of age-related neurodegenerative disorders.

### OGT^C921Y^ is defective in glycosyltransferase activity toward protein substrates in vitro

Disrupted OGT stability and folding caused by pathogenic missense variants could contribute to OGT-CDG pathophysiology (Pravata et al., 2020b). Therefore, the effect of C921Y substitution on the OGT protein structure was investigated. Although we were able to produce this variant in recombinant form from an *E. coli* expression system, we were unable to grow crystals for structural analysis, unlike several of the previously reported variants (Gundogdu et al., 2018; Pravata et al., 2019; Pravata et al., 2020a; Selvan et al., 2017; Vaidyanathan et al., 2017). Nevertheless, analysis of the wild type OGT structure reveals that C921 resides in the catalytic domain of OGT, proximal to the UDP-GlcNAc binding site (∼13 Å, Fig. 2A). The mutation of C921 to a bulky tyrosine residue would disrupt the C845-C921 disulfide bridge found in wild type OGT. Furthermore, C921Y could perturb the position of H911, which in the wild type enzyme establishes π−π stacking interactions with the UDP uracil ring (Fig. S2 A&B). The mutation could also affect N935, induce the loss of the Y851-UDP interaction and promote the generation of a new interaction between the Y931 and UDP (Fig. S2 A&B). Thus, the C921Y substitution could affect protein stability, interfere with catalysis and/or alter binding of UDP-GlcNAc to OGT.

**Figure 2:**
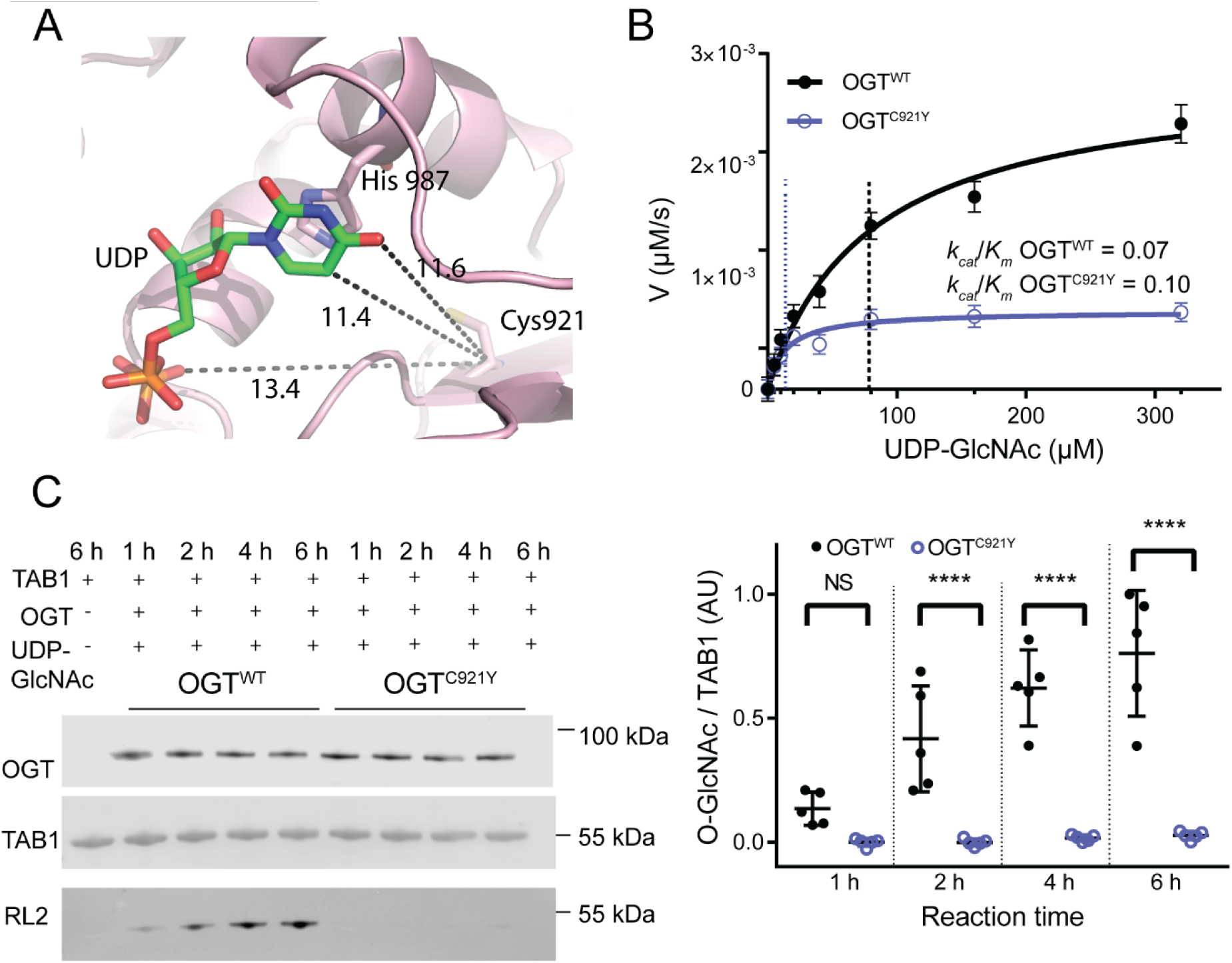
OGT^C921Y^ is defective in glycosyltransferase activity towards protein substrates *in vitro*. (a) Zoomed view into catalytic domain of OGT. UDP is shown as sticks. Distances between C921 residue and UDP are highlighted by dotted lines (13.3 Å from C921 to the uracil ring in green, 14.3 Å from C921 to the pyrophosphate linker in orange). (b) Michaelis-Menten kinetics of recombinant full length OGT^WT^ and OGT^C921Y^ incubated with acceptor peptide (Ac-KENSPAVTPVSTA-NH2) and varying concentrations of UDP-GlcNAc, n = 3 repeats, each consisting of three technical replicates. (c) Immunoblot detection of TAB1 O-GlcNAcylation by recombinant OGT^WT^ and OGT^C921Y^ (323-1041) over the course of 1 h, 2 h, 4 h and 6 h *in vitro*. Corresponding immunoblot signal quantification is shown on the right. n = 5 reactions using the same batch of recombinant protein. Repeated Measures two-way ANOVA with Sidak’s multiple comparison test, 0 h *p* = 0.14, 2 h *p* < 0.0001, 4 h *p* < 0.0001, 2 h *p* < 0.0001. Error bars represent standard deviation.

We next investigated whether the C921Y mutation disrupts stability of recombinant OGT (323 – 1044aa) *in vitro* using a thermal denaturation assay. No change in melting temperature between truncated OGT^WT^ (T_m_ = 45 ± 1 ⁰C) and OGT^C921Y^ (T_m_ = 45 ± 1 ⁰C) was observed (n = 3 replicates, each consisting of three technical repeats, Fig. S3). Next, we examined the impact of the C921Y mutation on the glycosyltransferase activity of full-length OGT. We first used a steady-state kinetics assay with varying concentrations of sugar donor UDP-GlcNAc against an established acceptor peptide (Ac-KENSPAVTPVSTA-NH_2_,(Pathak et al., 2015)) (Fig. 2B). The maximal reaction velocity (*V*_max_) and the Michaelis Menten constant (*K*_m_) were reduced for OGT^C921Y^ (*V*_max_ = 0.70 x 10^-3^ ± 0.01 µmol/s, *K*_m_ = 14 ± 4 µM) compared to OGT^WT^ (*V*_max_ = 2.8 x 10^-3^ ± 0.2 µmol/s, *K*_m_ = 78 ± 15 µM). However, the catalytic efficiency (k_cat_ / *K*_m_) of the mutant and wild type enzyme were similar (OGT^WT^ *k*_cat_ / *K*_m_ = 0.07, OGT^C921Y^ k_cat_ / *K*_m_ = 0.10, n = 3 replicates, each consisting of three technical repeats). Thus, the catalytic activity of OGT^C921Y^ towards a peptide substrate is intact compared to OGT^WT^.

We next evaluated effects of the mutation on OGT activity towards TAK1-binding protein 1 (TAB1), a well characterised protein substrate (Pathak et al., 2012; Rafie et al., 2017), revealing a loss of OGT^C921Y^ catalytic activity compared to OGT^WT^ (Fig. 2C). Western blot analysis of TAB1 O-GlcNAcylation showed that the time dependent increase in TAB1 O-GlcNAc signal produced by OGT^WT^ was absent in the OGT^C921Y^ catalyzed reaction (Fig. 2D). The mean level of TAB1 O-GlcNAcylation catalyzed by OGT^C921Y^ after 6 h was 28-fold lower than that catalyzed by OGT^WT^ (Two-way Anova with Sidak’s multiple comparison testing, *p* < 0.0001).

These data reveal that while OGT^C921Y^ catalytic activity towards protein substrates is abrogated, it appears intact towards peptides, implying that disruption of substrate interactions with the catalytic domain of OGT caused by the C921Y mutation prevents accommodation of large acceptor substrates. Interestingly, this suggests a previously unappreciated role of catalytic domain residues in recognition of protein substrate features beyond the site of O-GlcNAc modification. Together, these data show that OGT^C921Y^ is defective in glycosyltransferase activity toward protein substrates *in vitro*.

### OGT^C921Y^ disrupts O-GlcNAc homeostasis in undifferentiated mESCs

To investigate the impact of the OGT^C921Y^ variant on O-GlcNAc cycling, we engineered male OGT^C921Y^ mESC lines using CRISPR/Cas9. The murine and human OGT protein sequences are identical in the mutated region (Fig. S4A). Three independent CRISPR/Cas9 lines carrying OGT^C921Y^ were generated through clonal expansion from three distinct founder cells (Fig. S4B). OGT^C921Y^ mESC lines were morphologically identical to wild type cells and their cell cycle was not affected (Fig. S5). mRNA and protein levels of key pluripotency factors Oct4 and Sox2 remained the same as in OGT^WT^ mESC (Fig. S6). All the following experiments were performed using three different cell clones per genotype and repeated over multiple passages, unless stated otherwise.

Western blot analyses from pluripotent OGT^C921Y^ mESCs revealed a significant decrease in global O-GlcNAc levels compared to wild type cells (Figs. 3A,B, OGT^WT^ n = 15 biological replicates, OGT^C921Y^ n = 13 biological replicates, Unpaired t test, *p* = 0.0072), suggesting an alteration in O-GlcNAc homeostasis. We detected a significant upregulation of Ogt protein levels in OGT^C921Y^ cells compared to the wild type (Fig. 3A & C, OGT^WT^ n = 15 biological replicates, OGT^C921Y^ n = 13 biological replicates, Unpaired t test, *p* < 0.0001). However, given the decreased global O-GlcNAcylation levels in the mutant mESC lines, these data suggest that, despite being present at higher levels, OGT^C921Y^ glycosyltransferase activity is not sufficient to maintain O-GlcNAc homeostasis in mESCs. Oga protein levels were unchanged in the mutant cell line (Figs. 3A,D, OGT^WT^ n = 15 biological replicates, OGT^C921Y^ n = 13 biological replicates, Unpaired t test, *p* = 0.14), whereas previously characterized OGT-CDG variants showed a reduction in OGA protein levels when modelled in mESCs or patient-derived lymphoblastoids whilst OGT levels remained unchanged compared to wild type (Pravata et al., 2019; Pravata et al., 2020a; Vaidyanathan et al., 2017).

**Figure 3:**
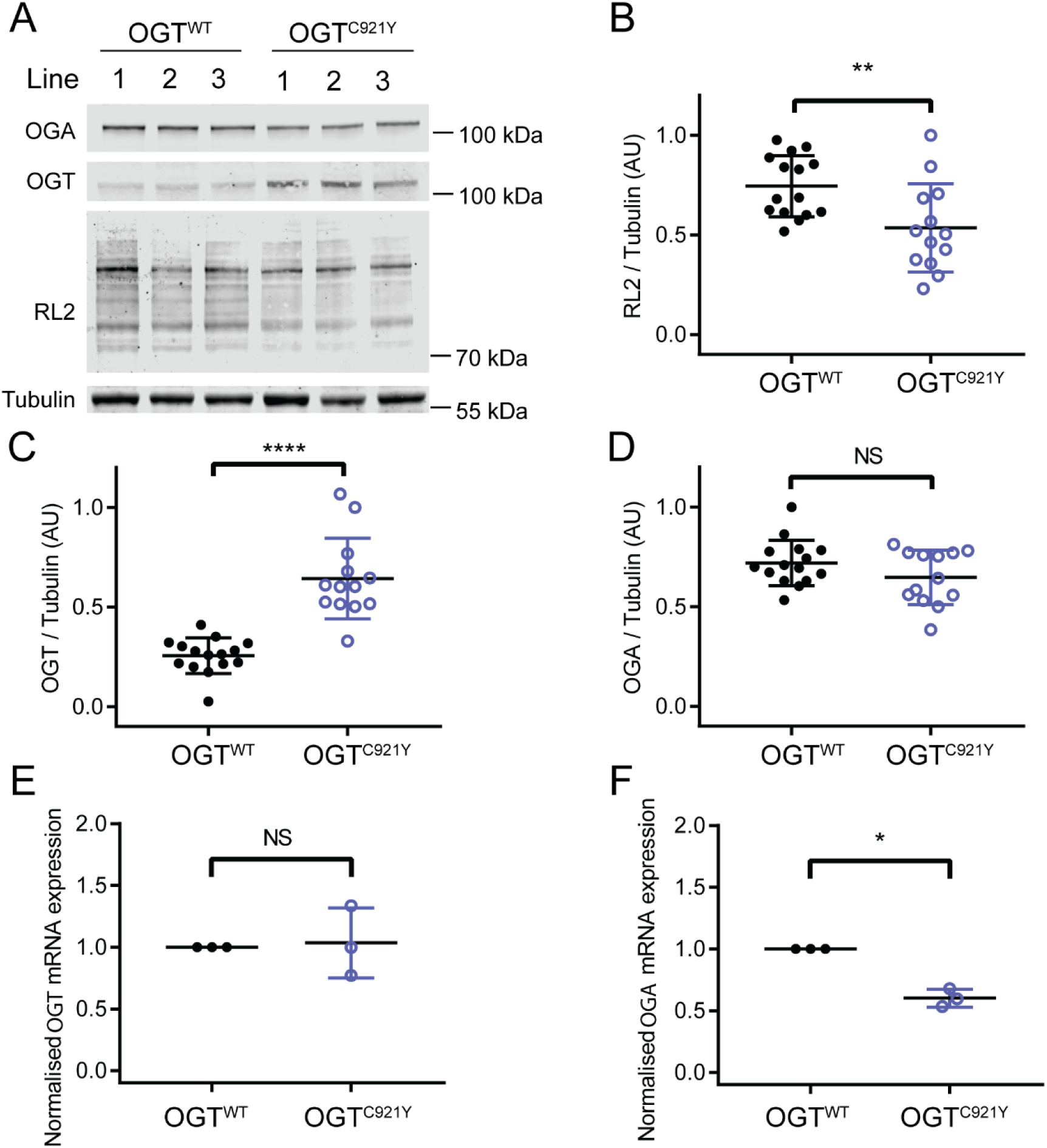
The OGT^C921Y^ variants disrupt O-GlcNAcylation machinery in undifferentiated mESCs. **(a)** Immunoblot showing detection of O-GlcNAcylation (RL2), OGT and OGA in mESCs carrying wild type OGT or the OGT^921Y^ variant. The quantification shown in panels B – D is based on results obtained from three different cell clones per genotype and repeated over four to five passages per clone. Error bars represent standard deviation. **(b)** Quantification of RL2 immunoblot signal. RL2 signal was normalized to Tubulin. OGT^WT^ n = 15, OGT^C921Y^ n = 13, Unpaired t test, *p* = 0.007. **(c)** Quantification of OGT immunoblot signal. OGT signal was normalized to Tubulin. OGT^WT^ n = 15, OGT^C921Y^ n = 13, Unpaired t test, *p* < 0.0001. **(d)** Quantification of OGA immunoblot signal. OGA signal was normalized to Tubulin. OGT^WT^ n = 15, OGT^C921Y^ n = 13, Unpaired t test, *p* = 0.14. **(e)** RT – PCR analysis of OGT mRNA normalized to GAPDH, 18 S and Actin β. Data points representing the mean expression calculated from three separate RT-PCR runs are shown. Each RT - PCR run was set up using several OGT^WT^ and OGT^C921Y^ as biological replicates. Unpaired *t* test, *p* = 0.84. **(f)** RT – PCR analysis of OGT mRNA normalized to GAPDH, 18 S and β-Actin. Data points representing the mean expression calculated from three separate RT-PCR runs are shown. Each RT-PCR run was set up using several OGT^WT^ and OGT^C921Y^ lines as biological replicates. Unpaired *t* test, *p* = 0.0007.

We next evaluated *Ogt* and *Oga* expression with RT-PCR analysis of three biological replicates with up to three OGT^C921Y^ and OGT^WT^ lines. These experiments showed that *Ogt* mRNA expression is unchanged in OGT^C921Y^ lines compared to wild type (Fig. 3E, n = 3 RT-PCR runs, each consisting of two to three biological replicates, Two-way Anova, *p* = 0.82), implying that Ogt may be stabilised at the protein level in OGT^C921Y^ mESCs, possibly through decreased protein degradation in cultured cells. Qian and colleagues (2018) revealed a bidirectional feedback mechanism between Ogt and Oga at the transcriptional level in primary mouse hepatocytes using an overexpression system. Furthermore, OGT has been shown to direct the transcriptional repressor Sin3A-HDAC1 complex to the *OGA* promoter in OGT-CDG patient-derived lymphoblastoids, thus modulating *OGA* expression (Vaidyanathan et al., 2017). Indeed, *Oga* mRNA levels were significantly decreased in OGT^C921Y^ mESCs (Fig. 3F, n = 3 RT-PCR runs, each consisting of two to three biological replicates, unpaired *t* test, *p* = 0.0007) despite protein levels being unchanged. Taken together, these data reveal that OGT^C921Y^ disrupts O-GlcNAc homeostasis in pluripotent mESCs despite compensatory changes in the levels of Ogt protein and *Oga* expression.

### HCF1 processing is unchanged in pluripotent OGT^C921Y^ mESCs

In addition to catalyzing O-GlcNAcylation, OGT also proteolytically processes and activates the transcriptional coregulator HCF1 (Capotosti et al., 2011). HCF1 is encoded by *HCFC1*, which itself has been reported to be an intellectual disability gene (Huang et al., 2012; Wongkittichote et al., 2021). Biochemical analysis of two previously reported OGT–CDG variants showed decreased HCF1 proteolysis (Pravata et al., 2019; Willems et al., 2017). HCF1 proteolysis occurs within the same active site of OGT as O-GlcNAcylation (Kapuria et al., 2016; Lazarus et al., 2013). Given the proximity of the C921Y substitution to the OGT active site (Figs. 2A, S1) and the impact of the C921Y variant on OGT glycosyltransferase activity, we investigated whether this variant leads to deficient HCF1 O-GlcNAcylation and processing. First, we performed an *in vitro* HCF1 repeat 1 (HCF1rep1) cleavage assay using full length recombinant hOGT^WT^ or hOGT^C921Y^ (Fig. S7). The uncleavable mutant HCF1rep1^E10D^ was used in this assay as a negative control to allow us to distinguish between unspecific HCF1 degradation and *bona fide* proteolytic products (Fig. S7). In agreement with the loss of O-GlcNAcylation activity on TAB1, we observed decreased levels of O-GlcNAcylation of HCF1rep1^WT^ and HCF1rep1^E10D^ by OGT^C921Y^ (Fig. 4A). However, both the wild type and mutant OGT were able to catalyse the formation of HCF1 proteolytic products (Fig. 4A). To corroborate this further, we investigated HCF1 cleavage in the mESC model (Fig. 4B). Since HCF1 translocates to the nucleus, the abundance of HCF1 proteolytic fragments was inspected in both the cytoplasmic and nuclear fractions of undifferentiated mESCs (Fig. 4B). We observed no difference in HCF1 signal between wild type OGT and OGT^C921Y^ cells in either of the cellular compartments (Fig. 4C, n = 6 biological replicates, One-way Anova with Tukey comparison test, cytoplasmic fraction *p* = 0.99, nuclear fraction *p* = 0.54). RT-PCR of HCF1 revealed that HCF1 mRNA levels remain stable in OGT^C921Y^ mESCs (Fig. 4D, n = 3 RT-PCR runs, each consisting of two to three biological replicates, unpaired *t* test, p value = 0.172). Taken together, these data show that HCF1 processing is unchanged in undifferentiated OGT^C921Y^ mESCs.

**Figure 4:**
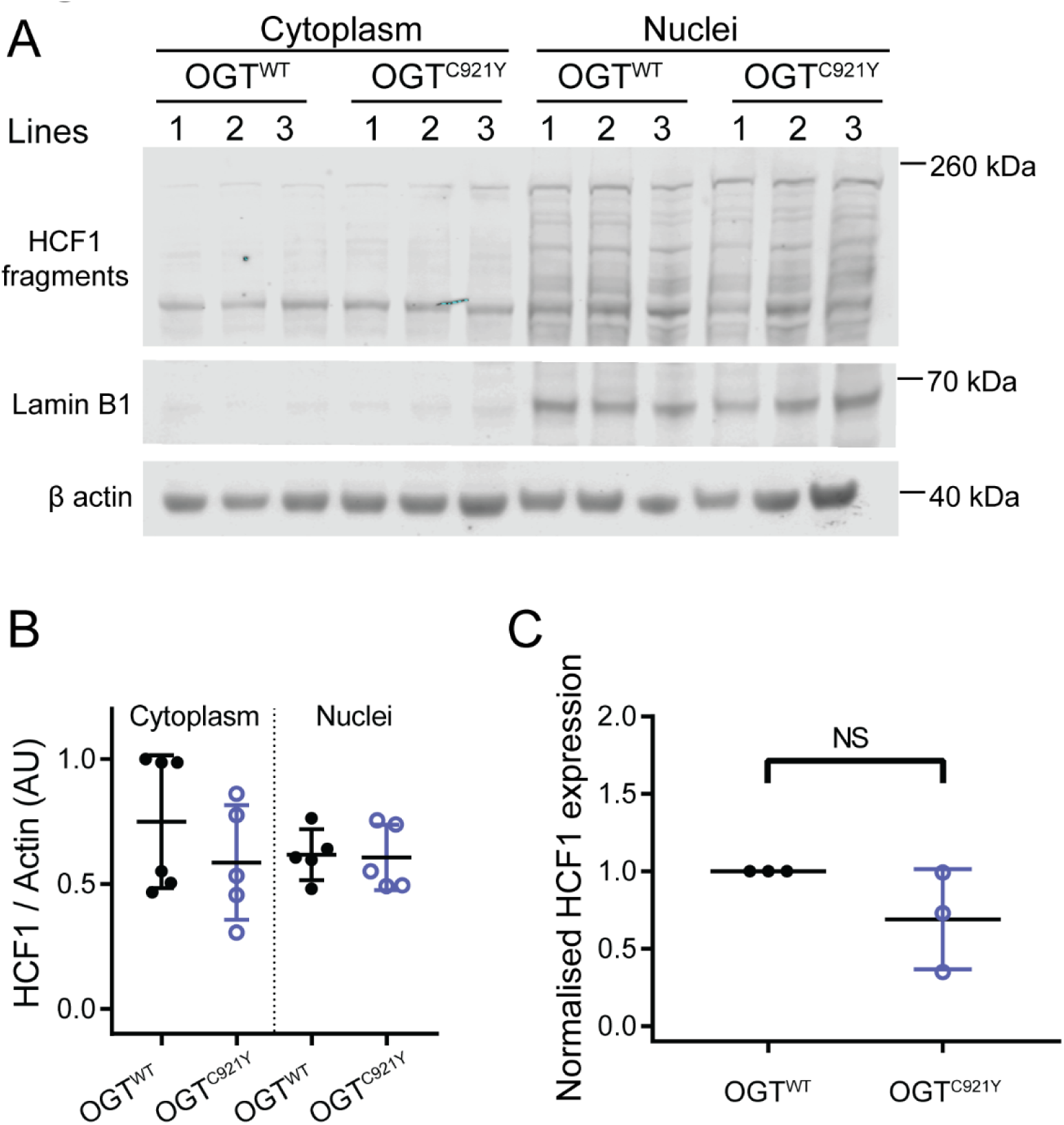
HCF1 processing by undifferentiated OGT^WT^ and OGT^C921Y^ in mESCs. **(a)** Immunoblot of HCF1 proteolytic fragments in the cytoplasm and nucleus of mESCs harboring either wild type or OGT^C921Y^ variant. Lamin B1 was used as marker for nuclear fraction. This experiment was performed using three cell clones per genotype and repeated over two passages. **(b)** Quantification of HCF1 signal. HCF1 signal was normalized to Actin β. n = 6 biological replicates, One-way Anova with Tukey comparison test, cytoplasmic fraction *p* = 1, nuclear fraction *p* = 0.54. Error bars represent standard deviation. **(c)** RT – PCR analysis of HCF1 mRNA expression normalized to GAPDH, 18 S and Actin β. Data points representing the mean expression calculated from three separate RT-PCR runs are shown. Each RT-PCR run was set up using several OGT^WT^ and OGT^C921Y^ lines as biological replicates. Unpaired *t* test, *p* = 0.172.

### OGT^C921Y^ abrogates mESC self-renewal capacity

Analogous to previously reported OGT-CDG mutations (Pravata et al., 2020b), patients carrying the OGT^C921Y^ variant present with a broad array of phenotypes including developmental delay, brain abnormalities and musculoskeletal defects. Embryonic development and patterning are crucially dependent on the ability of the inner cell mass of the early embryo to respond to the presence of differentiation stimuli or the absence of self-renewal stimuli. The ability of the inner cell mass as well as cultured ESCs to develop into the three primary germ cell layers is called pluripotency. Pluripotent cells maintain their identity through self-renewal. In the context of embryonic development, self-renewal and differentiation are meticulously orchestrated by various signalling molecules. In cell culture, mESCs are commonly propagated through LIF supplementation in the growth media (Smith et al., 1988; Williams et al., 1988). Upon LIF withdrawal, mESCs lose expression of pluripotency markers such as Oct4 and Nanog, colonies acquire a flattened morphology, and cells begin to differentiate (Chen et al., 2015; Cherepkova et al., 2016; He et al., 2017). O-GlcNAcylation has been previously shown to be important for core and auxiliary pluripotency factor function (Constable et al., 2017; Hao et al., 2019; Jang et al., 2012; Kim et al., 2021; Myers et al., 2016). Therefore, we hypothesized that the OGT^C921Y^ variant may impact the ability of stem cells to maintain an undifferentiated state. To test this hypothesis, we assayed colony formation and maintenance of stemness upon LIF withdrawal.

OGT^C921Y^ and wild type mESCs were subjected to clonogenic conditions in presence of LIF (positive control) or followed by LIF withdrawal for 24-96 h before fixing and alkaline phosphatase (ALP) staining (Fig. 5A). High expression of ALP is considered to be a marker of pluripotency, with ALP staining visualising pluripotent colonies (Štefková et al., 2015). The ALP-stained colonies were examined under a light microscope and classified into three categories (undifferentiated, mixed and differentiated) based on the intensity of ALP stain and colony morphology. Examples of colony scoring are shown in Fig. S8. In the wild type cell lines, the percentage of differentiated colonies increased after LIF withdrawal (Figs. 5B,C, n = 9 biological replicates, one-way Anova with Dunnett’s multiple comparison test, *p* < 0.0001) and the percentage of undifferentiated colonies significantly decreased with time of culture in absence of LIF (Figs. 5B,D, n = 9 biological replicates, one-way Anova with Dunnett’s multiple comparison test, *p* = 0.012). However, the percentages of undifferentiated and differentiated mutant OGT^C921Y^ colonies did not change significantly in relation to the length of LIF deprivation (Figs. 5B-D, n = 9 biological replicates, one-way Anova with Dunnett’s multiple comparison test, *p* = 0.10 and *p* = 0.29 respectively). We noted that OGT^C921Y^ mESCs produced significantly higher percentage of differentiated colonies than wild type mESCs at every LIF withdrawal time point apart from 96 h (Fig. 5B,C, n = 9 biological replicates, two-way Anova, 24 h, *p* < 0.0001, 48 h *p* = 0.02, 72 h *p* = 0.004, 96 h *p* = 0.13). Remarkably, OGT^C921Y^ colonies showed a significantly increased number of differentiated colonies even in presence of LIF (n = 9 biological replicates, two-way Anova, *p* = 0.0001) compared to wild type. The number of mixed colonies remained constant over the course of the assay in both wild type and OGT^C921Y^ mESCs. Furthermore, the overall number of colonies formed by OGT^C921Y^ mESCs was significantly lower than for wild type mESCs (Fig. S9, n = 45 wells scored, unpaired *t*-test, *p* < 0.0001). This observation can be explained by a decrease in clonogenic potential of OGT^C921Y^ compared to wild type, and/or increased cell death following cell plating at limiting density. Lastly, OGT^C921Y^ mESCs formed colonies with significantly larger surface area than wild type cells (Fig. S9, OGT^WT^ n = 2270 colonies measured, OGT^C921Y^ n = 1533 colonies measured, unpaired t test, *p* < 0.0001; observations were pooled together from all three independent experiments performed using three clones per genotype). Given the loss of ALP staining in OGT^C921Y^, the observed increase in surface area may reflect flattening and concomitant expansion of OGT^C921Y^ colonies due to differentiation.

**Figure 5:**
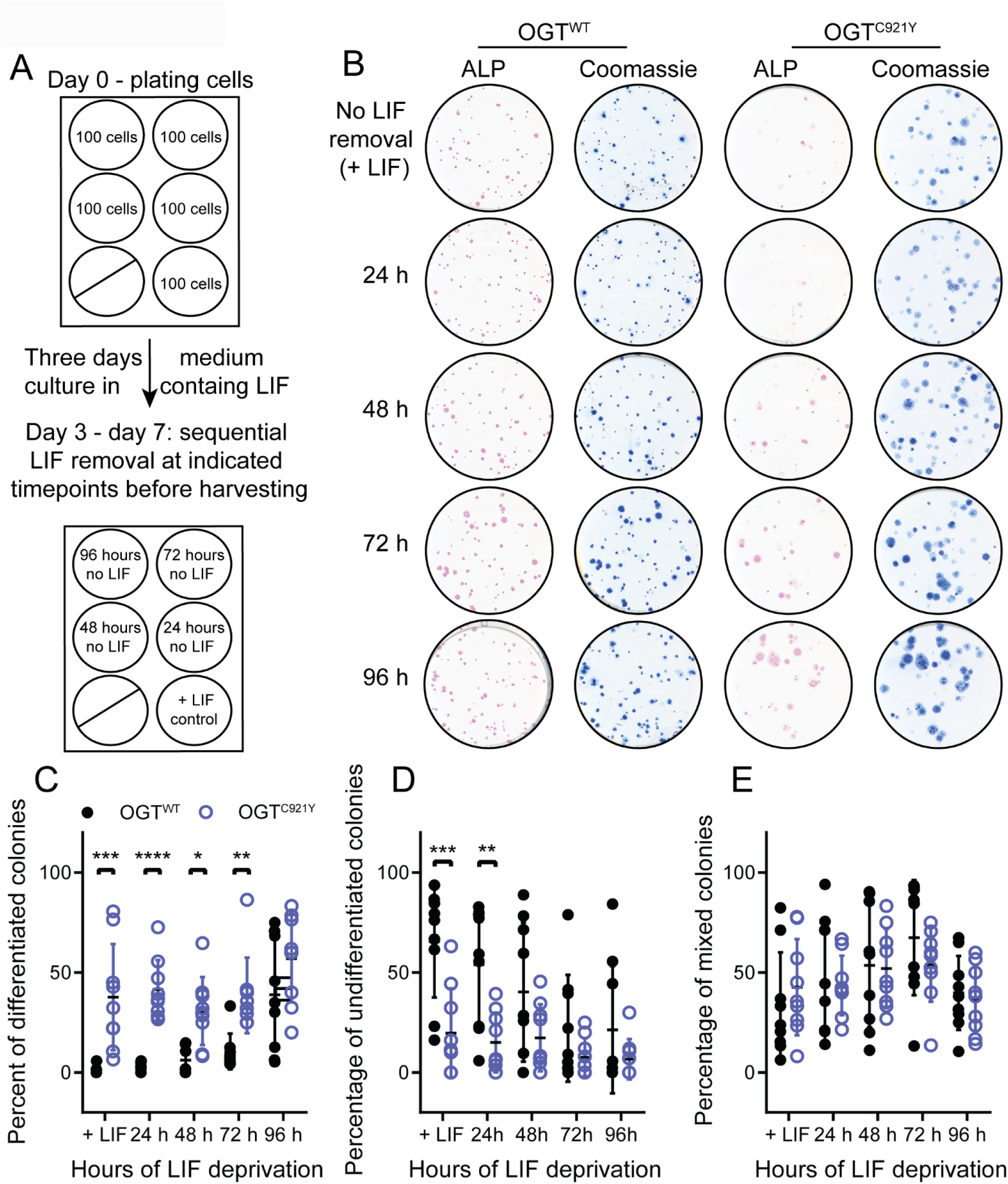
The OGT^C921Y^ variant leads to increased differentiation in response to LIF withdrawal in mESCs. **(a)** Schematic representation of experimental set up. This experiment has been performed using three cell clones per genotype and repeated over three passages. **(b)** Scans of alkaline phosphatase (ALP, pink) stained colonies of wild type or OGT^C921Y^ mESCs at different time points after LIF withdrawal. Each well was re-stained by Coomassie (blue) to reveal ALP negative colonies. **(c)** Plot of percentages of differentiated colonies in wild type and OGT^C921Y^ mESCs. n = 9 biological replicates, Two-way Anova, +LIF *p* = 0.0001 24 h *p* = 0.005, 48 h *p* = 0.21, 72 h *p* = 0.68, 96 h *p* = 0.67. **(d)** Plot of percentages of undifferentiated colonies in wild type and OGT^C921Y^ mESCs. n = 9 biological replicates, Two-way Anova, +LIF *p* = 0.0006, 24 h *p* < 0.0001, 48 h *p* = 0.017, 72 h *p* = 0.004, 96 h *p* = 0.13. **(e)** Plot of percentages of mixed colonies in wild type and OGT^C921Y^ mESC. n = 9 biological replicates, Two-way Anova, all *p* values are non-significant.

To further corroborate the O-GlcNAcylation status and enzymatic activity of OGT^C921Y^ mESCs under clonogenic conditions, we sought to investigate the levels of global protein O-GlcNAcylation and protein levels of key pluripotency transcription factors (Sox2 and Oct4). OGT^C921Y^ and OGT^WT^ mESCs were cultured for 7 days under clonogenic conditions in the presence of LIF (Fig. 6A). As opposed to the decreased glycosyltransferase activity of the OGT^C921Y^ variant *in vitro* and in confluent mESCs, the decrease in global O-GlcNAcylation in OGT^C921Y^ colonies compared to OGT^WT^ colonies was not significant (Fig. 6B,D, OGT^WT^ n = 8 biological replicates, OGT^C921Y^ n = 9 biological replicates, unpaired t test, *p* < 0.07) despite increased levels of the Ogt protein (Fig. 6B,C, n = 9 biological replicates, unpaired t test, *p* = 0.0004). Previous reports indicated that global O-GlcNAcylation is highest in pluripotent stem cells and decreases as pluripotency is lost (Liu et al., 2012; Sheikh et al., 2020). Our observation may therefore reflect the shift from predominantly pluripotent stem cell population present in confluent mESCs propagated in LIF to a mixed population of cells present at day six of the clonogenic assay (Fig. 5C-E). Furthermore, Western blot analysis of lysates derived from OGT^C921Y^ colonies revealed a significant decrease of Oct4 (Fig. 6B,G, n = 9 biological replicates, unpaired t test, *p* = 0.004) and Sox2 (Fig. 6B,F, n = 9 biological replicates, unpaired t test, *p* = 0.002) levels compared to OGT^WT^. This finding was corroborated by immunofluorescence staining of OGT^WT^ and OGT^C921Y^ colonies (Fig. S10). It is worth noting that we observed a variability in Sox2 and Oct4 protein levels among the three lines of OGT^C921Y^ mESCs, possibly stemming from the fact that each cell line is derived from a separate CRISPR/Cas9 event, therefore representing three cell populations that arose from three different founder cells. To discern whether differentiating OGT^C921Y^ mESCs assume specific germ layer preferentially, colonies were immunolabelled for markers specific for the mesoderm (Brachyury), endoderm (sox17) and ectoderm (pax6). There was no difference in germ layer marker protein expression in the mutant and wild type colonies (Fig. S11, S12, S13 and S14). However, this result is limited by the short timeframe of the assay. Taken together, these data show that independent of LIF signalling, OGT^C921Y^ mESCs show reduced ALP staining under clonogenic conditions compared to OGT^WT^, suggesting defects in the stem cell renewal. Furthermore, OGT^C921Y^ colonies grown in the presence of LIF show reduced levels of Sox2 and Oct4 transcription factors, implying that OGT^C921Y^ affects self-renewal of mESC under these conditions.

**Figure 6:**
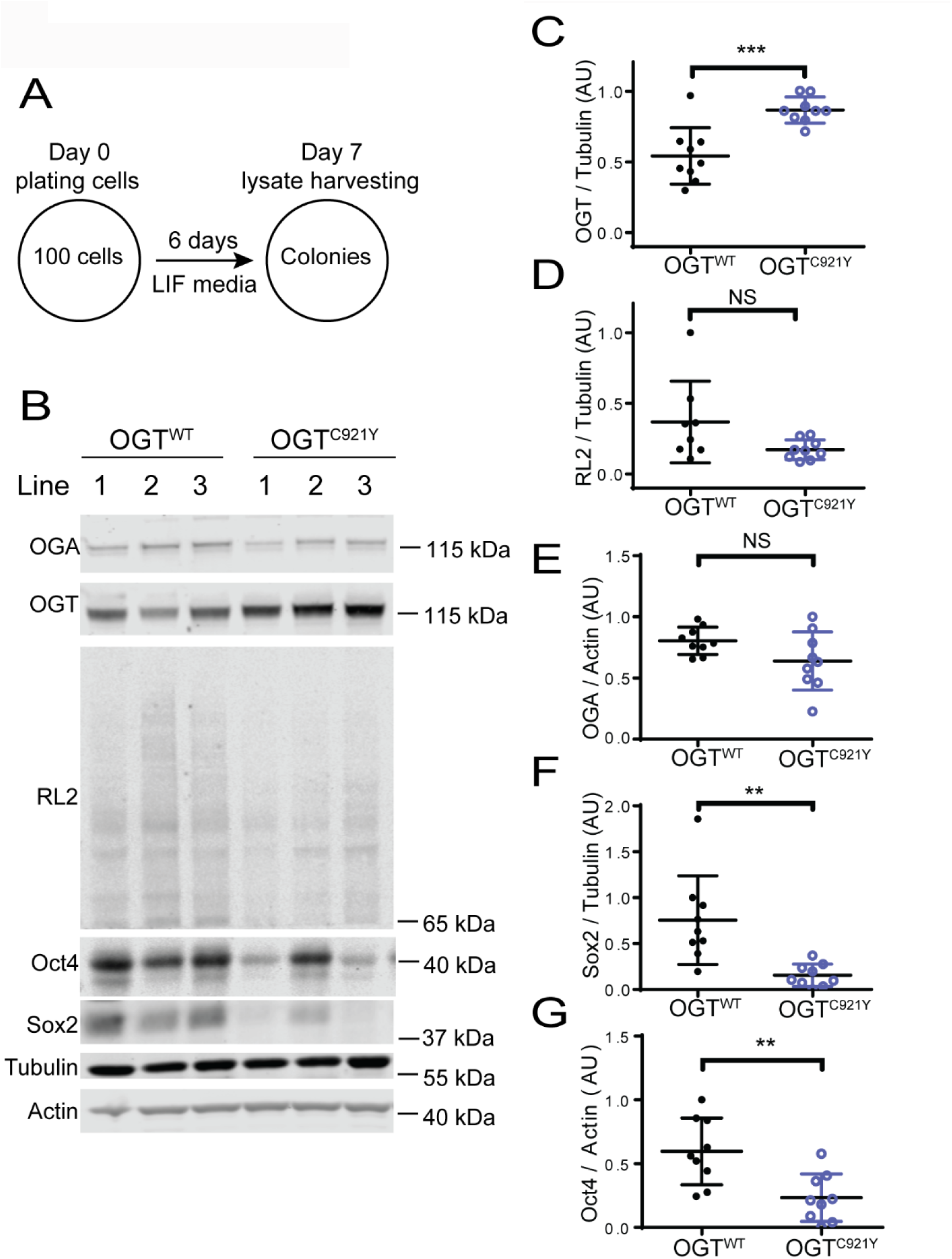
mESCs harboring the OGT^C921Y^ variant downregulate Sox2 and Oct4 in response to clonogenic conditions. **(a)** Schematic representation of experimental set up. The following experiments were performed using three cell clones per genotype and repeated over three passages. **(b)** Immunoblot showing detection of O-GlcNAcylation (RL2), OGT, OGA, Oct4 and Sox2 in mESC lines carrying wild type or OGT^921Y^ variant. Lysates were derived from cells grown under clonogenic conditions in presence of LIF as shown in (a). **(c)** Quantification of OGT immunoblot signal. RL2 signal was normalized to Tubulin. OGT^WT^ n = 9 biological replicates, OGT^C921Y^ n = 9 biological replicates, Unpaired t test, *p* = 0.0004. Error bars represent standard deviation. **(d)** Quantification of RL2 immunoblot signal. RL2 signal was normalized to Tubulin. OGT^WT^ n = 8 biological replicates, OGT^C921Y^ n = 9 biological replicates, Unpaired t test, *p* = 0.067. Error bars represent standard deviation. **(e)** Quantification of OGA immunoblot signal. OGA signal was normalized to Actin. OGT^WT^ n = 9 biological replicates, OGT^C921Y^ n = 9 biological replicates, Unpaired t test, *p* = 0.077. Error bars represent standard deviation. **(f)** Quantification of Sox2 immunoblot signal. Sox2 signal was normalized to Tubulin. OGT^WT^ n = 9 biological replicates, OGT^C921Y^ n = 9 biological replicates, Unpaired t test, *p* = 0.0023. Error bars represent standard deviation. **(g)** Quantification of Oct4 immunoblot signal. Oct4 signal was normalized to Actin. OGT^WT^ n = 9 biological replicates, OGT^C921Y^ n = 9 biological replicates, Unpaired t test, *p* = 0.0038. Error bars represent standard deviation.

### Concluding remarks

Pathogenic missense variants in the *OGT* gene co-segregate with intellectual disability in at least 17 known affected families, seven of which have been described in detail (Bouazzi et al., 2015; Pravata et al., 2019; Pravata et al., 2020a; Selvan et al., 2018; Vaidyanathan et al., 2014; Willems et al., 2017). Through biochemical, cellular, and animal model assays, recent studies revealed that these single *OGT* point mutations (hemizygous or heterozygous) underpin pathogenesis, *via* unknown biological mechanisms. This condition was recently classified as a distinct type of CDG, termed OGT-CDG (Pravata et al., 2020b). Here, we describe a family affected by a C921Y variant in the catalytic domain of OGT. Three male siblings suffering from ID harbour the OGT^C921Y^ variant, while their brother who possesses a wild type copy of *OGT* is healthy. All three affected brothers present with dysmorphic features such as long and pear-shaped nose, as well as behavioural and language problems. These features are in line with the recently reported general clinical phenotype description of OGT-CDG patients (Pravata et al., 2020b). The severity of the OGT^C921Y^ patients’ ID varied. Interestingly, the level of ID manifestations was different even between the two twins affected by this novel OGT variant. Similar differences in ID severity were reported by Pravata and colleagues in female monozygotic twins suffering from OGT-CDG (Pravata et al., 2019).

The C921Y mutation is predicted to cause structural defects within the OGT catalytic core, albeit without measurable effects on stability *in vitro*. Interestingly, only two of the OGT-CDG variants biochemically characterized to date (OGT^N567K^ and OGT^N648Y^) did not lead to changes in OGT stability (Pravata et al., 2019; Pravata et al., 2020a). OGT^N567K^, OGT^N648Y^ and OGT^C921Y^ are the only three described variants directly affecting the catalytic domain, whereas other OGT-CDG mutations reside in the TPR domain and result in destabilization of the protein. Collectively, these data suggest that either loss of functional OGT protein or disruption of OGT catalytic activity lead to the same clinical phenotype. Biochemical characterization of further variants affecting catalysis, solution of their crystal structures and examination of OGT localization in OGT-CDG cells will contribute to testing this hypothesis.

Previous research has revealed a role of the OGT TPR domain in protein - protein interactions and substrate recognition (Iyer and Hart, 2003; Jínek et al., 2004; Lazarus et al., 2013; Levine et al., 2018; Pathak et al., 2015; Rafie et al., 2017), demonstrating the role of an asparagine ladder in substrate binding. Our data imply a potential role of catalytic domain residues in substrate recognition, substantiated by predicted structural changes in the proximity of the active site and a loss of glycosyltransferase activity towards a protein substrate, without loss of activity towards a peptide substrate. We also revealed that Ogt activity in cultured mESCs was affected by the C921Y variant, leading to global hypo O-GlcNAcylation. Decreased O-GlcNAcylation in cultured cells as a result of an OGT-CDG mutation has previously only been observed in the OGT^N648Y^ variant (Pravata et al., 2020a). O-GlcNAc homeostasis is tightly regulated by feedback loops between OGT, OGA and O-GlcNAc levels (Decourcelle et al., 2020; Lin et al., 2021; Muthusamy et al., 2015; Tan et al., 2021; Zhang et al., 2014). This has been shown to occur through both translational and post-translational mechanism, for example, through regulation of intron detention, which alters the abundance of productive transcripts of *OGT* and *OGA* (Tan et al., 2021). Unlike in *Ogt^C921Y^* cells, this generally manifests as downregulation of OGA at the protein level, particularly upon OGT inhibition or in OGT-CDG catalytic domain mutations (Ortiz-Meoz et al., 2015; Pravata et al., 2019; Pravata et al., 2020a). However, reducing

O-GlcNAcylation through inhibiting the hexosamine biosynthetic pathway has been shown to increase OGT protein levels (Lin et al., 2021). Importantly, OGA is also implicated in neurodevelopment (Olivier-Van Stichelen et al., 2017) and cognitive functioning (Muha et al., 2020), both in model animals and potentially in humans (as determined by a genome wide association study on intelligence) (Savage et al., 2018). These data together suggested the hypothesis that reduction of OGA levels in itself could be a potential mechanism that underpins OGT-CDG (Pravata et al., 2020b). However, unlike other characterized OGT-CDG mESC cell lines, the OGT^C921Y^ variant did not induce OGA downregulation in mESCs. Furthermore, based on Western blot analysis, HCF1 proteolytic processing and localization was not affected in cultured mESCs in our study, even though O-GlcNAcylation and HCF1 proteolytic processing occur within the same active site of OGT. A previously reported OGT-CDG mutation affecting the TPR domain results in a similar phenotype: unaffected HCF1 processing and reduced glycosyltransferase activity (Gundogdu et al., 2018; Vaidyanathan et al., 2017). These data point to loss of O-GlcNAcylation on specific proteins as a likely link between OGT-CDG missense mutations and patient phenotypes.

To determine whether OGT-CDG may arise due to decreased O-GlcNAcylation in stem cells, we assayed self-renewal and differentiation in mESCs harbouring the C921Y mutation in *Ogt*. In these cells, the pluripotency marker ALP and the key pluripotency transcription factors Oct4 and Sox2 are significantly decreased in the presence of LIF, compared to wild type colonies, implying a defect in the maintenance of an undifferentiated state. The colonies formed by OGT^C921Y^ mESCs were flat and spread-out, resembling differentiating colonies in their morphology. In addition, global O-GlcNAcylation in OGT^C921Y^ mESC colonies is significantly reduced compared to OGT^WT^ mESC colonies. This is in line with a previous study indicating that preventing O-GlcNAcylation blocks mESC self-renewal (Jang et al., 2012). O-GlcNAcylation has also been shown to decrease during development (Liu et al., 2012), further supporting the hypothesis that decreased O-GlcNAcylation may be pathogenic through reducing stem cell pluripotency and self-renewal. However, the OGT^C921Y^ variant does not result in preferential differentiation towards a specific germ layer, despite previous research indicating that O-GlcNAcylation of SOX2 can promote differentiation towards an ectodermal lineage (Kim *et al.,* 2021). These findings suggest that a potential contributor to OGT-CDG clinical manifestation is a misregulation of exit from a pluripotent state in stem cells. This may occur at several stages of development, with recent evidence suggesting normal O-GlcNAcylation is required for maintenance not only of ESCs (Jang *et al.,* 2012), but also neural stem cells (NSCs) (White *et al.,* 2020, Shen *et al.,* 2021).

## Materials & Methods

### Sequencing of patient genetic material

The inclusion of patients in research studies has been approved by the local ethical committee and consent for publication of data and photographs was given by the family. Whole exome sequencing was performed on patient III:2. Subsequent screening for the OGT-variant was done by Sanger sequencing in the mother, the healthy brother (III:1) and the affected twins (III:3 and III:4). All genetic analysis was done at an accredited clinical laboratory in Amplexa Genetics A/S, Sverigesgade 24, Odense C, Denmark.

### Protein expression and purification

Human OGT^WT^ and OGT^C921Y^ constructs (full length and a 323 – 1044aa shortened construct with truncated TPR domain fused with N-terminal His tag (full length) and GST tag (323 – 1044 OGT)) were expressed in *E. coli* BL21 cells in plasmids carrying an ampicillin resistance cassette. Transformed colonies that incorporated the plasmids were selected on ampicillin agar plates, expanded overnight at 37 ⁰C in a shaking incubator in 5 x ampicillin LB broth as a starter culture, and then grown in desired quantity (6 – 12 liters) at 37 ⁰C until reaching 0.5 – 0.6 OD_600_. Subsequently, the temperature of the shaking incubator was decreased to 18 ⁰C, cultures were induced with 100 µM IPTG and further cultured for 16 h. Following overnight culture, bacteria were pelleted for 45 min at 4 ⁰C, resuspended in base buffer (0.1 M Tris-HCl, pH 7.5, 0.15 M NaCl, 0.5 mM TCEP) and lysed using a French press in the presence of 0.1 mg/ml DNase I, 0.5 mg/ml lysozyme and protease inhibitor cocktail (1 mM benzamidine, 0.2 mM PMSF, 5 mM leupeptin). Resulting lysates were then pelleted at 33 000 rcf at 4 ⁰C for 45 min. Supernatant was filtered with a 0.2 µM filter and the pellet was discarded. Clarified supernatant was exposed either to glutathione Sepharose 4B beads for GST tagged protein or NiNTA resin for His tagged protein for affinity purification. Bound protein was either eluted using 50 mM glutathione (for GST tagged protein) or 300 mM imidazole (for His tagged protein) in base buffer, buffer exchanged into base buffer only and the tag was cleaved off using 80 units of PreScission protease per prep. Protein was further purified using size exclusion chromatography and stored in solution with 25 % glycerol at – 80 ⁰C.

### Enzyme assays

Michaelis Menten kinetics of full length recombinant OGT^WT^ and OGT^C921Y^ against acceptor peptide (Ac-KENSPAVTPVSTA-NH_2_) were tested in a fluorometric *in vitro* assay as described by Pathak and colleagues (Rafie et al., 2017). The reaction time was 4 h at room temperature. Experiments were repeated three times on different days, each repeat consisting of three technical replicates. Glycosyltransferase activity of recombinant OGT^WT^ and OGT^C921Y^ (323 – 1044) against protein substrate was tested *in vitro* using TAB1 (7 – 420) as described previously (Rafie et al., 2017). Experiments were repeated on five times on different days.

### Differential scanning fluorimetry (DSF) assay

To prepare assay mixtures, OGT^WT^ and OGT^C921Y^ (323 – 1044) were diluted to 1.2 µM in base buffer containing 50 mM HEPES/NaOH pH 7.5, 150 mM NaCl, 0.5 mM TCEP and mixed with 1:5000 SYPRO Orange Protein Gel Stain (Sigma). 50 μL of assay mixture was dispensed per well in a white bottom qPCR plate and each condition was performed in technical triplicate. The experiment was repeated three times on different days. The prepared assay plate was exposed to temperature increases from 25 ⁰C to 95 ⁰C with 1 ⁰C increments for 5 s each using a CFX Connect Real-Time PCR Detection System (BioRad). SYBR fluorescence was detected after every temperature increase. Data were truncated using Excel and analysed in Prism (GraphPad) as described previously (Huynh and Partch, 2016).

### Cloning and CRISPR/Cas9 generation

Generation of OGT^C921Y^ mESCs was performed as described previously by (Pravata et al., 2019) with mutation specific parameters. A repair template was generated to create the C921Y mutation. This was cloned as a *Bam*HI-*Not*I fragment into a plasmid based on pGEX6P1. Paired gRNA sequences were selected and cloned as annealing oligos into pBABED-U6 and pX335-U6 plasmids. Silent mutations to remove the gRNA recognition sequences, in addition to a change at codon 935, were introduced by PCR of a gene block followed by restrictionless cloning into the cloned genomic region. The change at position 935 was introduced by the use of the C921Y wobble primer along with the reverse screening primer to generate a mutagenic PCR product and introduced by restrictionless cloning. The repair template was confirmed by DNA sequencing. Sequences of the listed custom-made reagents are listed in Supplementary Table 1.

Clones were screened using paired screening primers. These generated a 600bp PCR product which was then digested with Fastdigest *Xce*I. Wild type clones show two fragments – 277bp and 323bp. Successful incorporation of the repair template results in loss of restriction of the PCR product. Intact PCR products were confirmed by DNA sequencing. Following DNA confirmation of several clones, RNA was extracted, and one-step RT-PCR was carried out using Takara Primescript High fidelity RT-PCR kit with mOGT_solid_ex11_fwd and mOGT_solid_end_rev primers. The resulting PCR product was sequenced to confirm that the region outwith the repair template had not been damaged during repair and mRNA expression included the change.

### Tissue culture

AW2 mESC culture was performed as described previously in (Pravata et al., 2019). AW2 line is derived from E14-TG2a.IV (129/Ola) ES cells kindly donated from the MRC Centre for Regenerative Medicine, Institute for Stem Cell Research, University of Edinburgh (Zhou et al., 2013). Cells were routinely tested for mycoplasma and kept in strictly sterile conditions with no visible bacterial or fungal infection.

### Western blotting

Protein was extracted from confluent mESCs. Plates or flasks with attached cells were washed twice with pre-warmed 1 x PBS and then covered with 20 µl per cm^2^ with ice cold lysis buffer (50 mM Tris pH 7.4, 1 mM EDTA-NaOH, 1 mM EGTA-NaOH, 1% Triton X-100, 1 mM Na_3_VO_4_, 50 mM NaF, 0.27 M Sucrose, 5 mM protease inhibitor cocktail). Then, prepared plates were frozen at – 20 ⁰C overnight. Cells were then scraped off the plates, vortexed for 10 s and centrifuged for 45 min at 15 000 rcf at 4 ⁰C. Protein concentration in the clarified lysate was determined using Pierce™ 660 nm Protein Assay Reagent (Thermo). For sample resolution and protein detection, 20 µg of protein was mixed with LDS buffer, boiled, loaded onto a 4 – 12 % Bis – Tris gel (Invitrogen) and then transferred onto 0.2 µm nitrocellulose membrane (GE Healthcare). Following membrane blocking with 5 % BSA in 1 x TBS, primary antibodies were applied: Anti-O-GlcNAc Transferase (DM-17, Sigma-Aldrich, catalogue number O6264, 1:5000), RL2 (Novus, catalogue number NB300, 1:1000), O-GlcNAcase (1:500, Sigma, catalogue number SAB420026), Oct3/4 (C-10, Santa Cruz Biotechnologies, catalogue number sc-5279, 1:500), Sox2 (Santa Cruz Biotechnologies, catalogue number sc-365823, 1:500), Histone 3 (Cell Signaling Technologies, catalogue number 9715, 1:2000) and α-Tubulin (Proteintech, catalogue number 11224-1-AP, 1:5000). After incubation with the corresponding LI-COR secondary antibodies (1:10 000), signal was detected using a LI-COR Odyssey scanning system and quantified using Image Studio Lite (LI-COR). Data were normalized and analyzed in Prism (GraphPad). Number of repeats per experiment is detailed in the figure legends.

For HCF1 cleavage detection, nuclear fractionation was performed. Briefly, confluent cells were detached from plates using accutase, spun down and washed. Pelleted cells were then covered with ice cold buffer A (10 mM HEPES pH 7.5, 1.5 mM MgCl_2_, 10 mM KCl, 0.5 mM DTT, 0.05% NP40 and protease inhibitor cocktail), vortexed and centrifuged at 3000 rpm for 10 min at 4 °C. The resulting supernatant was used as the cytoplasmic lysate fraction. Pelleted material was treated with ice cold buffer B (5 mM HEPES pH 7.9, 1.5 mM MgCl_2_, 0.2 mM EDTA, 0.5 mM DTT and 26% glycerol (v/v)), homogenized on ice and centrifuged at full speed for 10 min at 4°C. The resulting supernatant was used as the nuclear lysate fraction. Due to low protein concentration, protein was precipitated from samples as described previously by (Clark et al., 2013). 20 µg of protein was then loaded onto 4 – 12 % gel as described above. Primary antibodies used were HCF1 (Bethyl, 1:1000), Laminin B1 (ZL-5, Abcam, 1:5000) and β-Actin (Proteintech, 1: 10 000). After incubation with corresponding LI-COR secondary antibody (1:10 000), signal was detected using a LI-COR Odyssey scanning system and quantified using Image Studio Lite (LI-COR). Data were normalized and analyzed in Prism (GraphPad).

### RT - PCR

RNA was extracted from confluent cells using the RNeasy kit (Qiagen). RNA purity was determined using Nanodrop 1000 (Thermo) and RNA concentration was measured using the Qubit^TM^ RNA Broad Range kit (Thermo). RNA was transcribed into cDNA using the qScript cDNA Synthesis Kit (Quantabio). qPCR reactions were set up in 384 well plates in a total volume of 10 µl using PerfeCTa® SYBR® Green FastMix® for iQ™ (Quanta) with 5 ng of cDNA and 250 nM forward + reverse primer mix. Three housekeeping genes (GAPDH, 18 S and Actin β) were used as reference genes and each condition was run in technical triplicate. No template as well as no reverse transcription controls were used. Reactions were run in a BioRad CFX 384 real time detection system (BioRad). Primer sequences are summarized in Supplementary Table 2. Results were analysed using BioRad CFX Manager and Prism (GraphPad). Number of repeats per experiment is detailed in the figure legends.

### Alkaline phosphatase assay

For alkaline phosphatase (ALP) assays, cells were well resuspended, passed through a cell strainer to remove clumps and doublets and counted using a hemocytometer. Then, 100 wild type or OGT^C921Y^ mESCs were plated into five wells of a 6-well plate pre-treated with 0.1 % (w/v) porcine gelatin in complete GMEM as described above. Cells were grown in complete media for two days to start forming colonies. From day three until day seven, media was changed in dedicated wells into no-LIF GMEM (GMEM (Gibco) supplemented with 5 % FBS fraction V (Gibco), 1 x NEAA (Gibco), 1 x sodium pyruvate (Gibco), 0.1 mM BME (Thermo)). One well per plate remained without media change. On day seven, media was aspirated, colonies were washed twice with 1 x PBS and fixed with 1 % paraformaldehyde for 2 min. Fixed colonies were stained with the Alkaline Phosphatase kit (Merck) according to manufacturer’s instructions. ALP-treated plates were scanned on a flat-bed scanner, and colonies counted and scored (undifferentiated, mixed, differentiated) under a microscope. Colonies were subsequently re-stained with Coomassie to visualize ALP-negative colonies and plates re-scanned. Experiment has been repeated three times using three cell lines per genotype. Colony diameters were determined using ImageJ based on Coomassie scans using a macro attached in the supplementary material. Briefly, colonies were segmented by creating a binary image by setting an automatic threshold using the “MaxEntropy” algorithm and measured using the “Analyze Particles” function. Macro is provided in supplementary materials. Data were analysed using Prism (GraphPad).

### Cell cycle analysis

Cell cycle analysis was performed using near-confluent OGT^C921Y^ (two lines) and OGT^WT^ (one line) mESCs collected over three subsequent passages. DNA content was assessed using Abcam Propidium Iodide kit (ab139418) according to manufacturer’s instructions. Flow cytometry analysis was performed on BD FACSCanto (BD). Data were analysed using FlowJo and Prism (GraphPad) software.

### Immunofluorescence analysis of colonies

Colonies were plated on coverslips coated with 0.1% (w/v) porcine gelatin in complete GMEM as described above. 6 days after plating, colonies were fixed for 15 min using 4% PFA. PFA was then quenched with 1 x TBS for 10 min. Fixed colonies were then washed with 1 x PBS twice for 5 min each. Samples were subsequently permeabilized and precipitated using ice cold 90% (w/w) methanol at -20 ℃ for 10 min. Samples were then washed twice with 1 x PBS and blocked blocking buffer (1 x PBS, 1% Tween20 and 5% BSA) for at least 15 min. Primary antibodies (Oct3/4 (D6C8T, Cell Signaling Technology, catalogue number 83932, 1:1000), Sox2 (D9B8N, Cell Signaling Technology, catalogue number 23064, 1:1000), Pax6 (H-295, Santa Cruz, catalogue number sc-11357, 1:1000), Sox17 (Santa Cruz, catalogue number sc-130259, 1:1000) and Brachyury (D10, Santa Cruz, catalogue number sc-166962, 1:1000)) were added in blocking buffer overnight. Following primary antibody stain, cells were washed three times for 15 min with 1 x PBS and secondary antibodies (1:2000) were added in blocking buffer. Colonies were washed with 1 x PBS and DAPI, mounted and imaged on a confocal microscope. Images were analyzed using an ImageJ macro, attached in the supplementary material. Briefly, nuclear regions were determined by performing a series of global and local thresholding steps. Non-nuclear regions were determined based on subtraction of nuclear regions from a binary image created by applying an automatic threshold to individual channels using the “MinError” algorithm.

## Acknowledgements

This work was funded by a Wellcome Trust Investigator Award (WT110061) to D.M.F.v.A. M.O. was funded by a Wellcome Trust 4-year PhD studentship (WT102132). I.C. was funded by a studentship from the National Centre for the Replacement, Refinement and Reduction of Animals in Research (NC3Rs, T001682). We would like to thank the patients and their family who consented to participate in this study. We acknowledge Dr. Lindsay Davidson for advice on experiment design.

## Author contributions

M.O., T.B.H., Ch.D.F. and D.M.F.v.A conceived the study; A.B. and T.B.H. performed clinical assessment, Ch.D.F. performed the exome analysis, M.O. performed experiments; M.M. and M.O. performed microscopy and cell cycle experiments, I.C. wrote code for alkaline phosphatase and microscopy images analysis, S.G.B. performed molecular docking simulation, V.M.P. generated the CRISPR cell line; A.T.F. performed molecular biology; M.O., M.P.S., A.B. and D.M.F.v.A. interpreted the data and wrote the manuscript with input from all authors.

## Supplementary information

**Supplementary figure S1:** *Figure removed for BioRxiv submission.* Profile photos and close up view of patient’s eyebrows. a) and b) Profile photographs of affected brothers carrying OGT^C921Y^ variant. c) Proband III.2 has sparse eyebrows.

**Supplementary figure S2:**
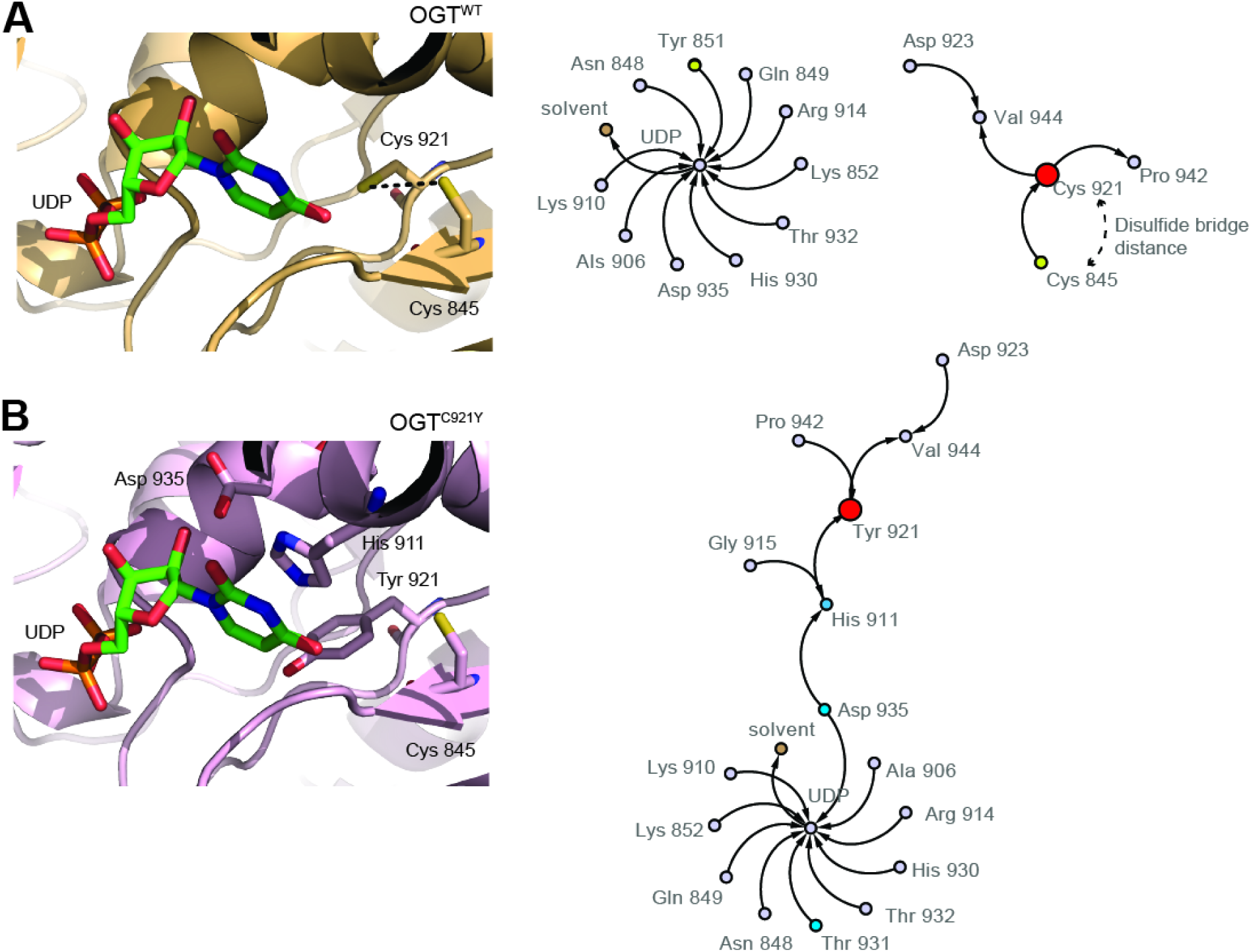
Model of the OGT C921Y substitution and changes in interactions in the catalytic core. a) The PDB 5LWV containing a UDP molecule sitting in the active site (green sticks) and a fusion protein peptide, removed for our analysis, was used a template for the OGT^WT^ protein. Averaged MD trajectory calculated after 1000 poses is shown in the left side. For clarity, cysteine 921 is represented as a red circle. b) The OGT^C921Y^ variant was modelled over the PDB 5LWV with the same modifications as described for OGT^WT^. Averaged MD trajectory calculated after 1000 poses is shown in the left side. For clarity the mutant tyrosine 921 is represented with a red circle. All key residues are labelled.

**Supplementary figure S3:**
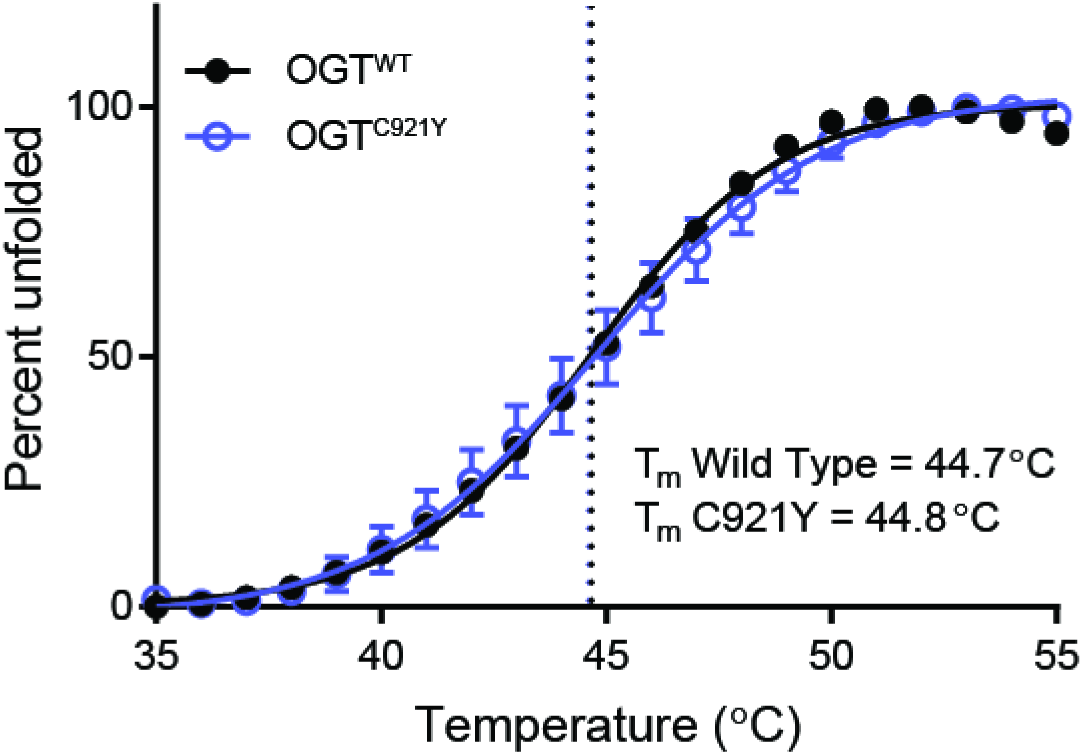
Melting temperature determination for OGT^WT^ and OGT^C921Y^ (323 – 1044). Thermal shift assay performed with recombinant OGT^WT^ and OGT^C921Y^ hOGT (323-1041). This experiment was performed three times, using three technical replicates each time. Error bars represent standard deviation.

**Supplementary figure S4:**
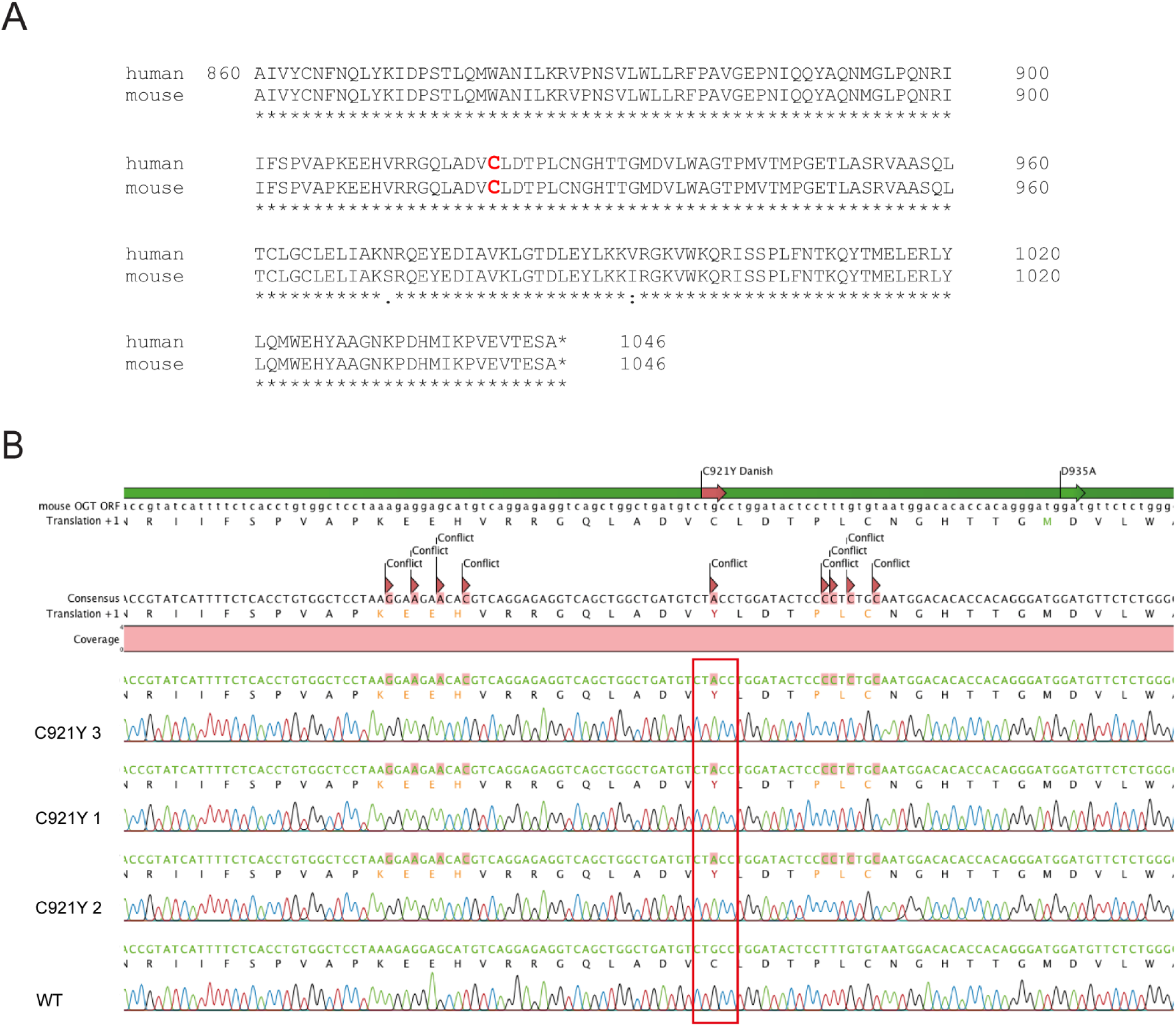
Generation of OGT^C921Y^ mESC lines. a) Sequence alignment of human and mouse OGT (amino acids 840 – 1046). Cysteine 921 is highlighted in red. b) DNA sequencing of generated OGTC921Y mESC clones (chromas 1 – 3) and wild type mESC (Chroma 4). Positions corresponding to cysteine 921 are highlighted in red rectangle.

**Supplementary figure S5:**
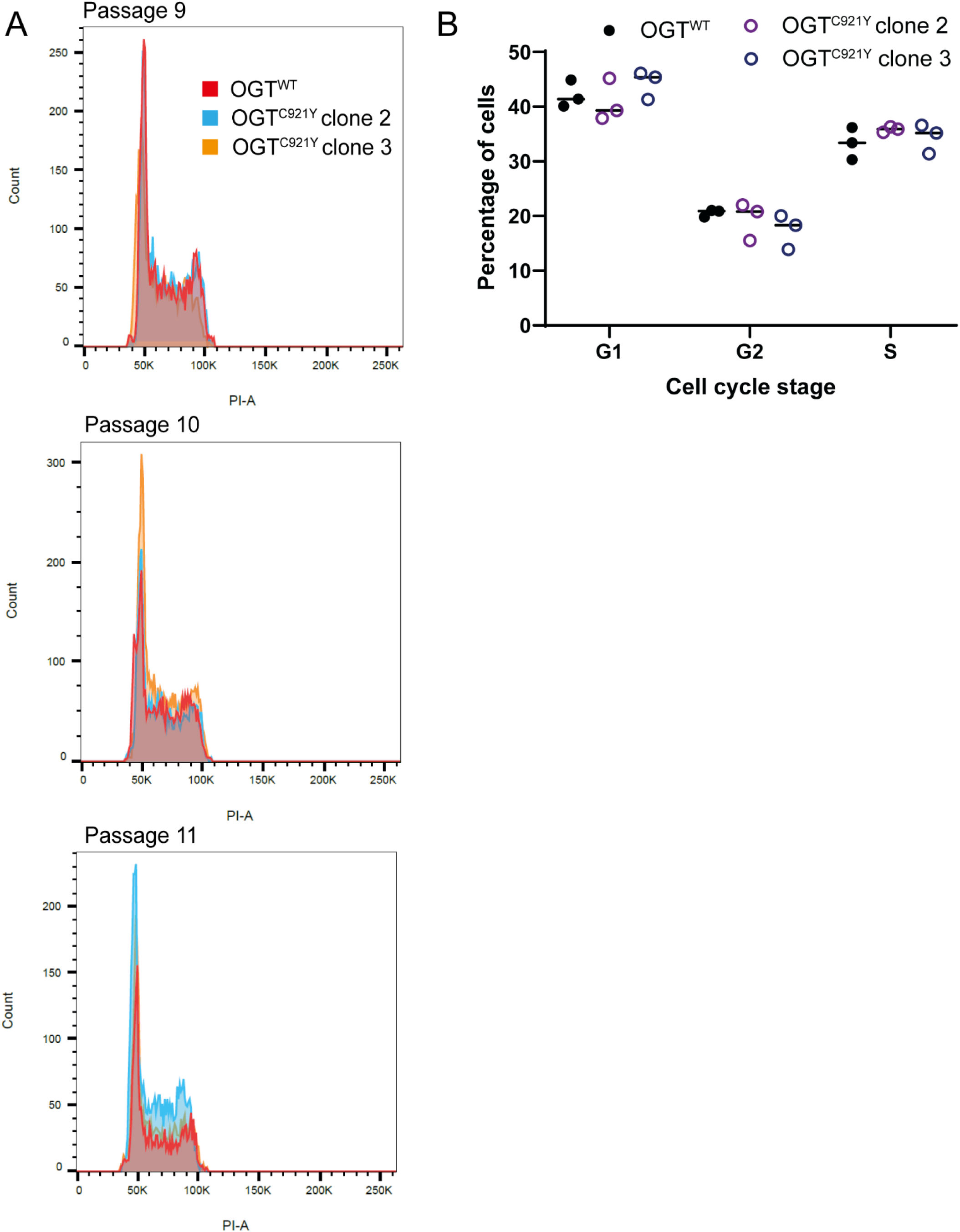
Cell cycle analysis of OGT^WT^ and OGT^C921Y^ mESC. a) Cell cycle profiles of OGT^WT^ and OGT^C921Y^ across three passages using two mutant and one wild type cell line. We aimed to perform cell cycle profiling with the earliest available passages, which is the reason for not using all three cell OGT^C921Y^ cell lines in this experiment. b) Plot of percentages of cells in each cell cycle stage.

**Supplementary figure S6:**
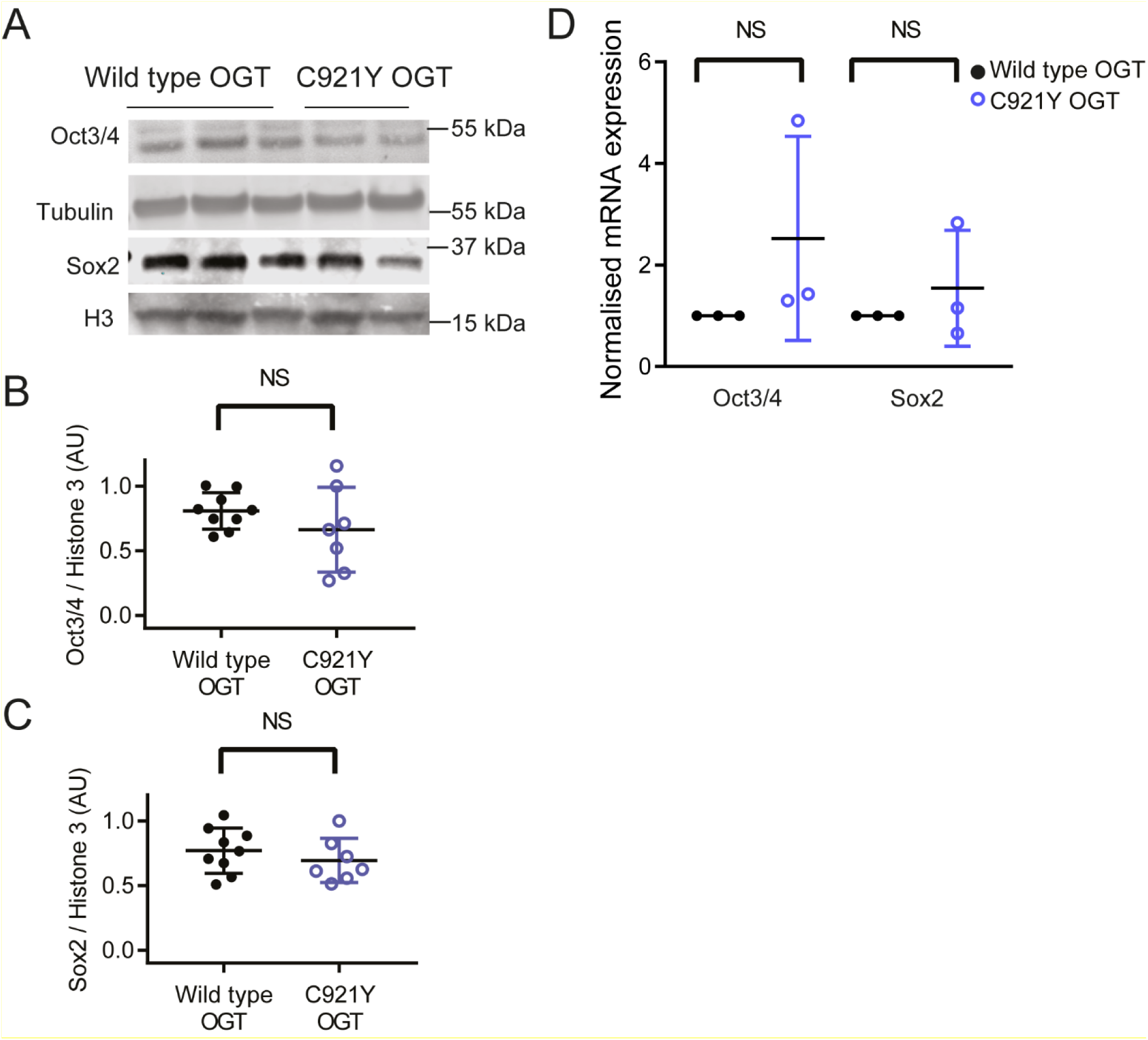
Oct3/4 and Sox2 expression in OGT^WT^ and OGT^C921Y^ mESC propagated in LIF. The quantification shown in panels B – D is based on results obtained from two to three different cell clones per genotype and repeated over three passages per clone. a) Immunoblot of Oct3/4, Sox2, Tubulin and Histone3. b) Quantification of Oct3/4 signal relative to tubulin. Unpaired *t* test, *p* value = 0.251. Error bars represent standard deviation. c) Quantification of Sox2 signal relative to Histone 3. Unpaired *t* test, *p* value = 0.401. Error bars represent standard deviation. d) mRNA expression of Oct3/4 and Sox2. Unpaired *t* test, Sox2 *p* value = 0.455, Oct3/4 *p* value = 0.260. Each data point represents normalised mean expression calculated from three separate RT-PCR runs. Each RT - PCR run was set up using several OGTWT and OGTC921Y as biological replicates. Error bars represent standard deviation.

**Supplementary figure S7:**
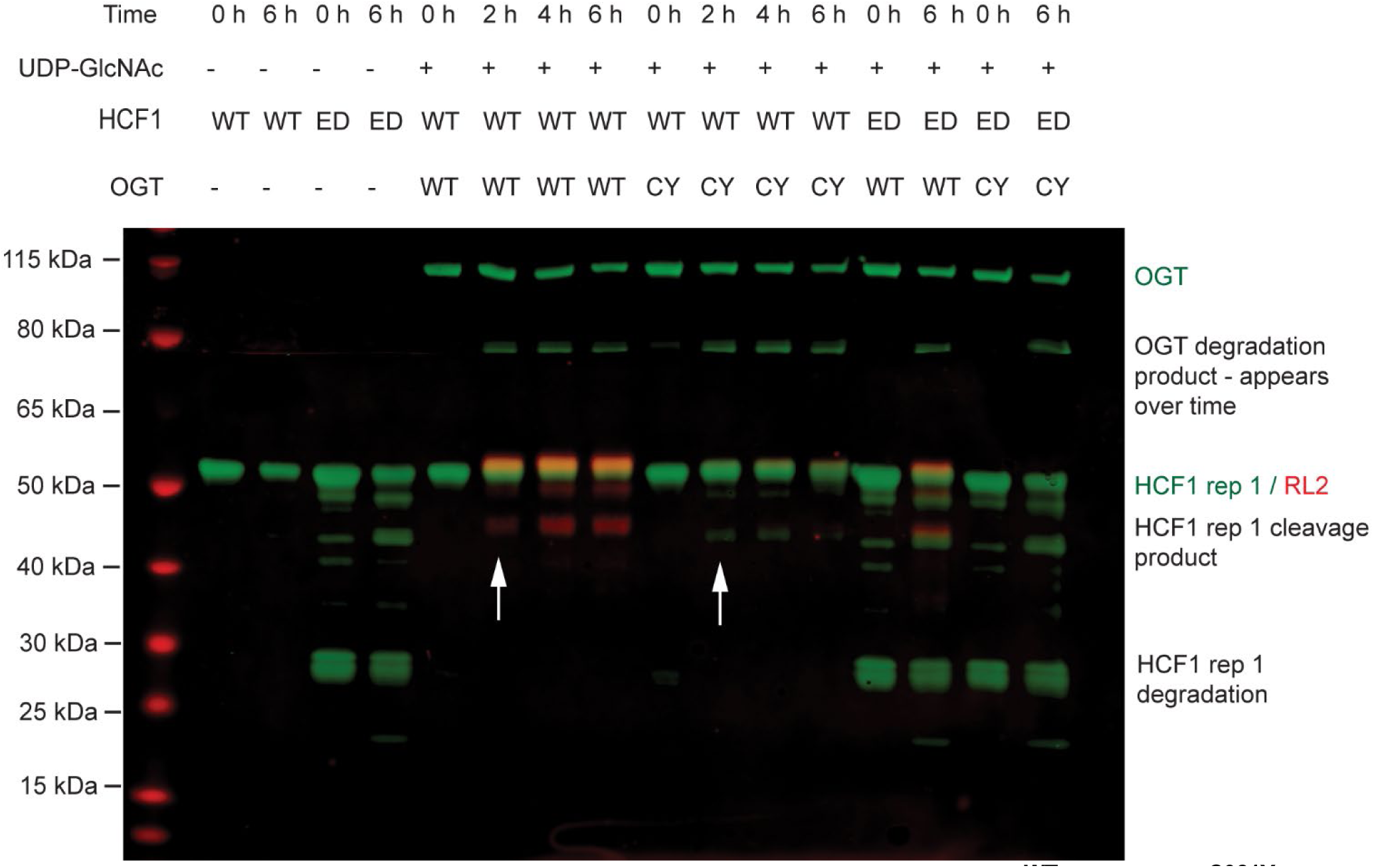
HCF1 processing by recombinant OGT^WT^ and OGT^C921Y^. Immunoblot of HCF1 proteolytic fragments produced in a time course *in vitro* reaction using the following human recombinant proteins: wild type (WT) and uncleavable (ED) HCF1 repeat 1 (HCF1rep1); OGT^WT^ and OGT^C921Y^. HCF1 and OGT were detected in the green channel, O-GlcNAc (RL2) was detected in the red channel.

**Supplementary figure S8:**
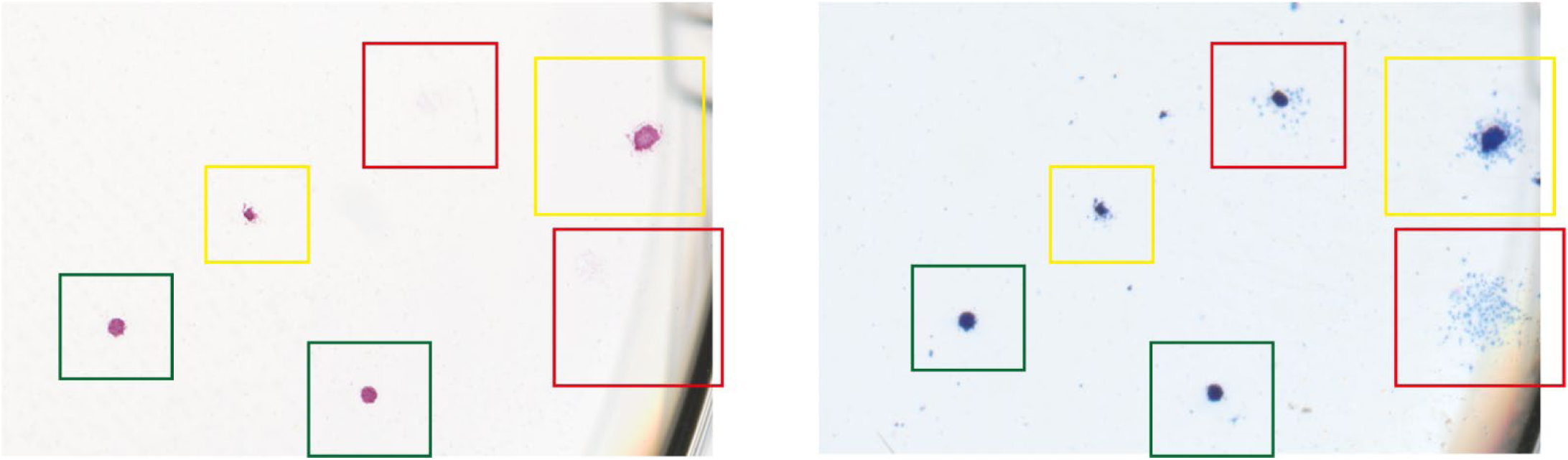
Colony scoring criteria. Images corresponding to the same section of a 6-well plate are shown. Image in the left represents colonies stained with ALP and the image on the right represents colonies stained with Coomassie. Compact colonies with strong ALP staining and no signs of differentiation were scored as undifferentiated (green rectangle), ALP positive colonies with signs of differentiation around the perimeter were scored as mixed (yellow rectangle) and dispersed colonies with no ALP signal were scored as differentiated (red rectangle).

**Supplementary figure S9:**
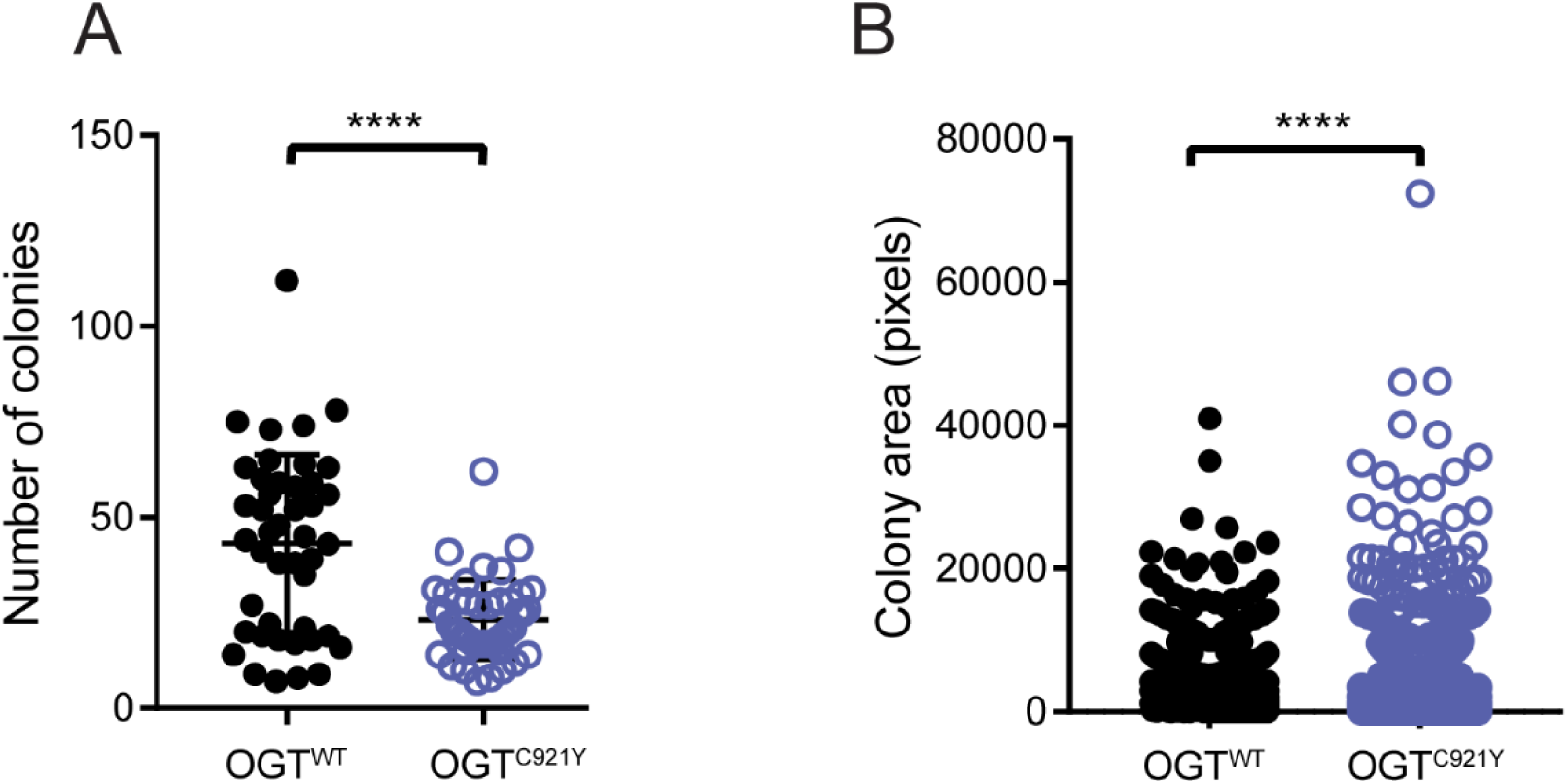
Number and size of OGT^WT^ and OGT^C921Y^ colonies. (a) Total number of colonies produced by wild type or OGT^C921Y^ mESCs in any condition. Unpaired t test, n = 45 wells scored, *p* < 0.0001. Error bars represent standard deviation. (b) Mean colony area with SEM measured by automated ImageJ macro based on scans of colonies stained with Coomassie. Unpaired t test, wild type n = 2270 measured colonies, OGT^C921Y^ n = 1533 measured colonies, *p* < 0.0001.

**Supplementary figure S10:**
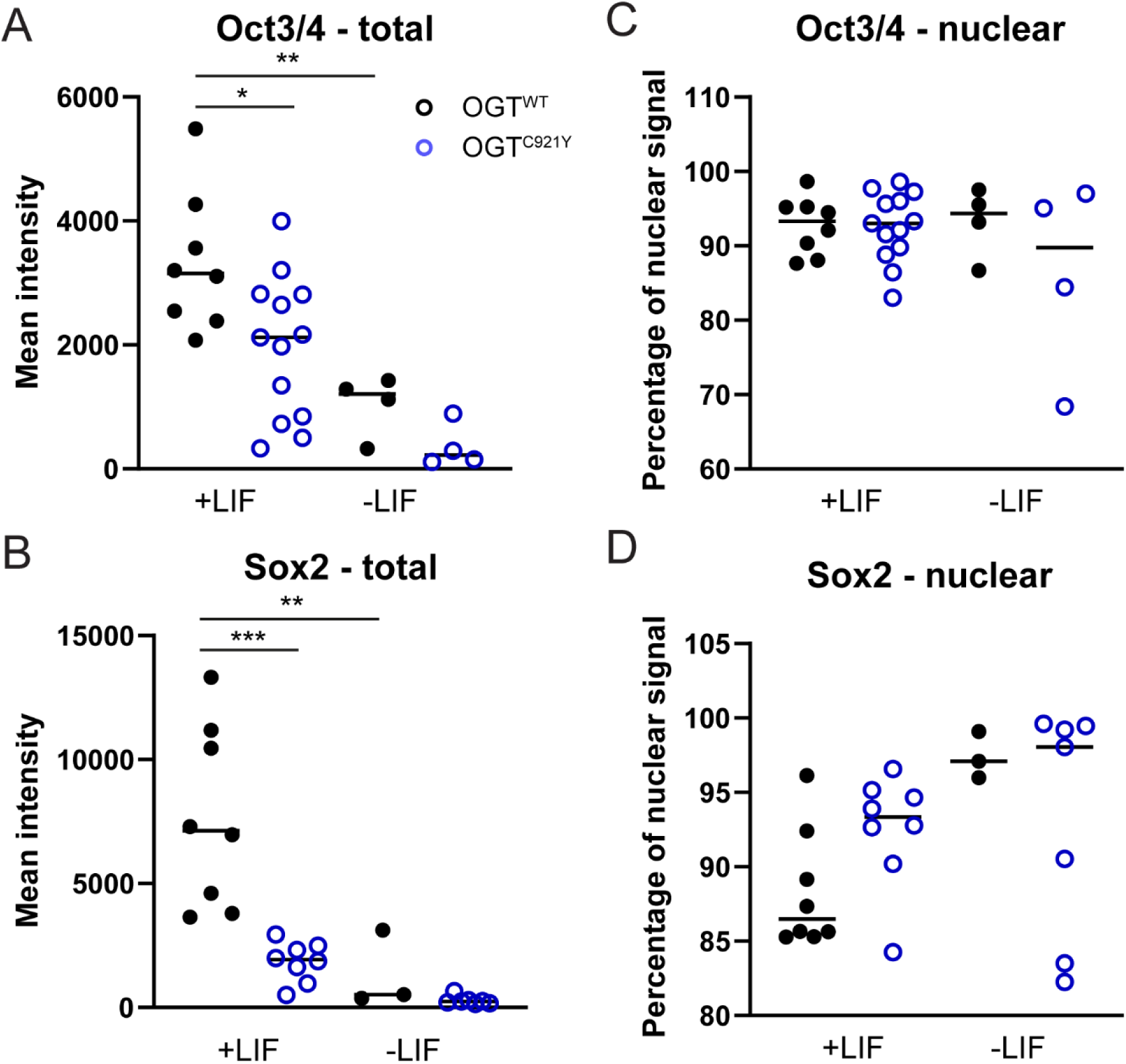
Oct3/4 and Sox2 levels in OGT^WT^ and OGT^C921Y^ colonies. (A, B) Each data point represents the mean intensity per colony calculated as sum signal intensity divided by colony area. Oct3/4 +LIF OGT^WT^ versus OGT^C921Y^, *p* value = 0.035 (One-way ANOVA followed by Sidak’s multiple comparison test). Oct3/4 +LIF OGT^WT^ versus -LIF OGT^C921Y^, *p* value = 0.006 (One-way ANOVA followed by Sidak’s multiple comparison test). Sox2 +LIF OGT^WT^ versus OGT^C921Y^, *p* value = 0.0001 (One-way ANOVA followed by Sidak’s multiple comparison test). Sox2 +LIF OGT^WT^ versus -LIF OGT^WT^, *p* value = 0.002 (One-way ANOVA followed by Sidak’s multiple comparison test). (D – F) Each data point represents the percentage of nuclear signal calculated as sum nuclear signal divided by sum signal intensity. Two lines per genotype over two passages were used for this experiment. All the colonies present on a coverslip were imaged. One-way ANOVA showed no significant differences in nuclear signal between tested groups.

**Supplementary figure S11:**
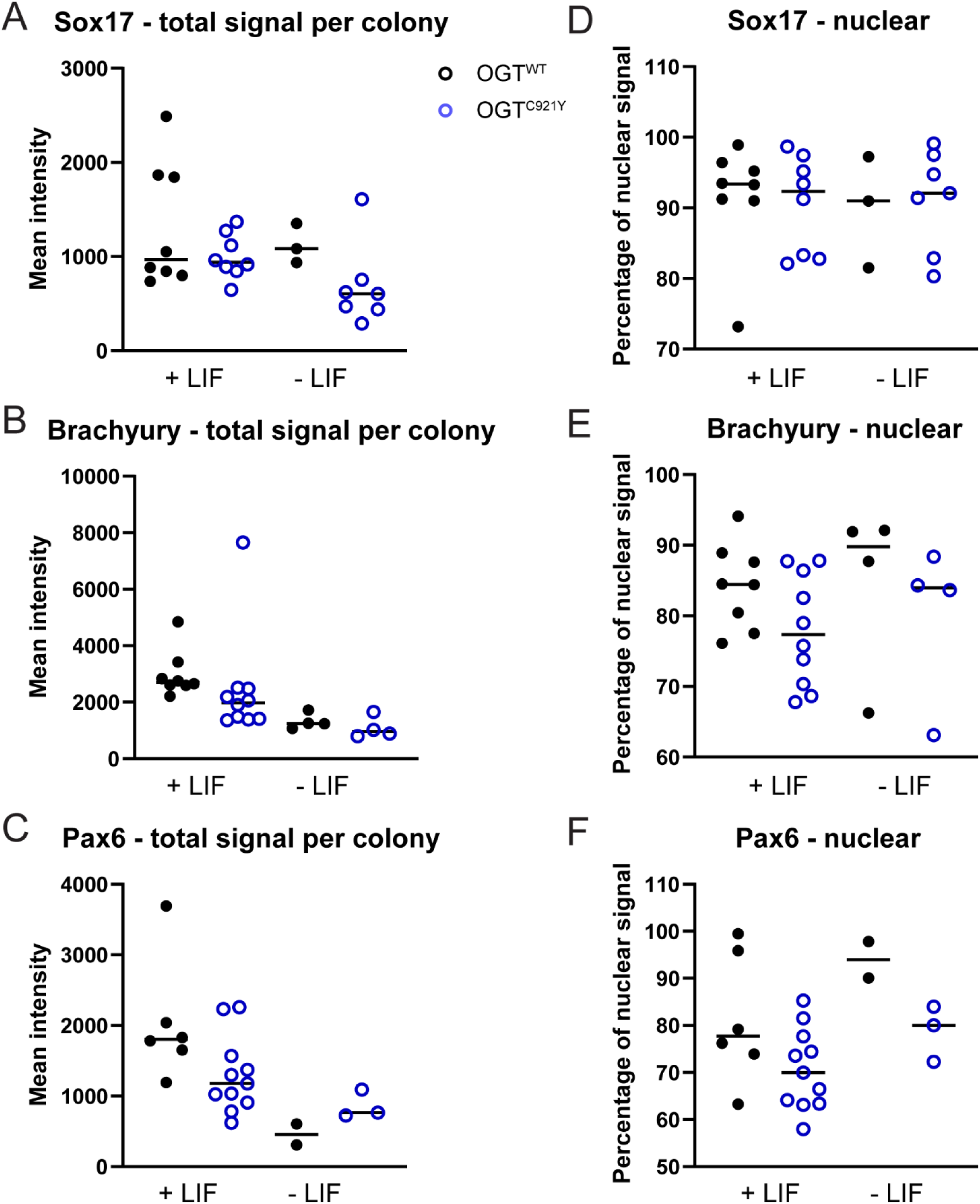
Sox17, Brachyury and Pax6 levels in OGT^WT^ and OGT^C921Y^ colonies. (A – C) Each data point represents the mean intensity per colony calculated as sum signal intensity divided by colony area. (D – F) Each data point represents the percentage of nuclear signal calculated as sum nuclear signal divided by sum signal intensity. Two lines per genotype over two passages were used for this experiment. All the colonies present on a coverslip were imaged. One-way ANOVA showed no significant differences between tested groups.

**Supplementary figure S12:**
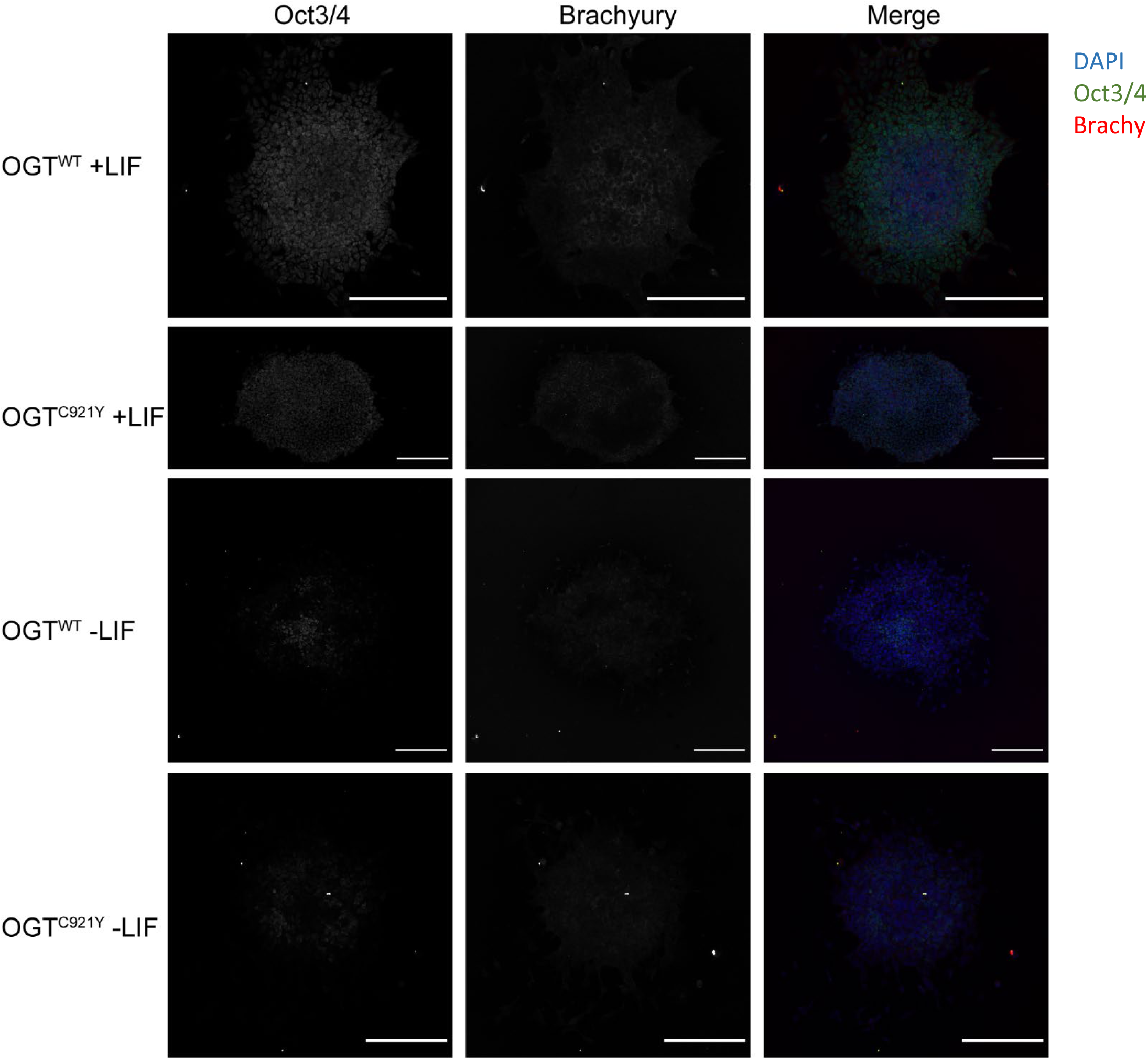
epresentative images of Oct3/4 and Brachyury staining in OGT^WT^ and OGT^C921Y^ colonies. Scale bar = 200 µM.

**Supplementary figure S13:**
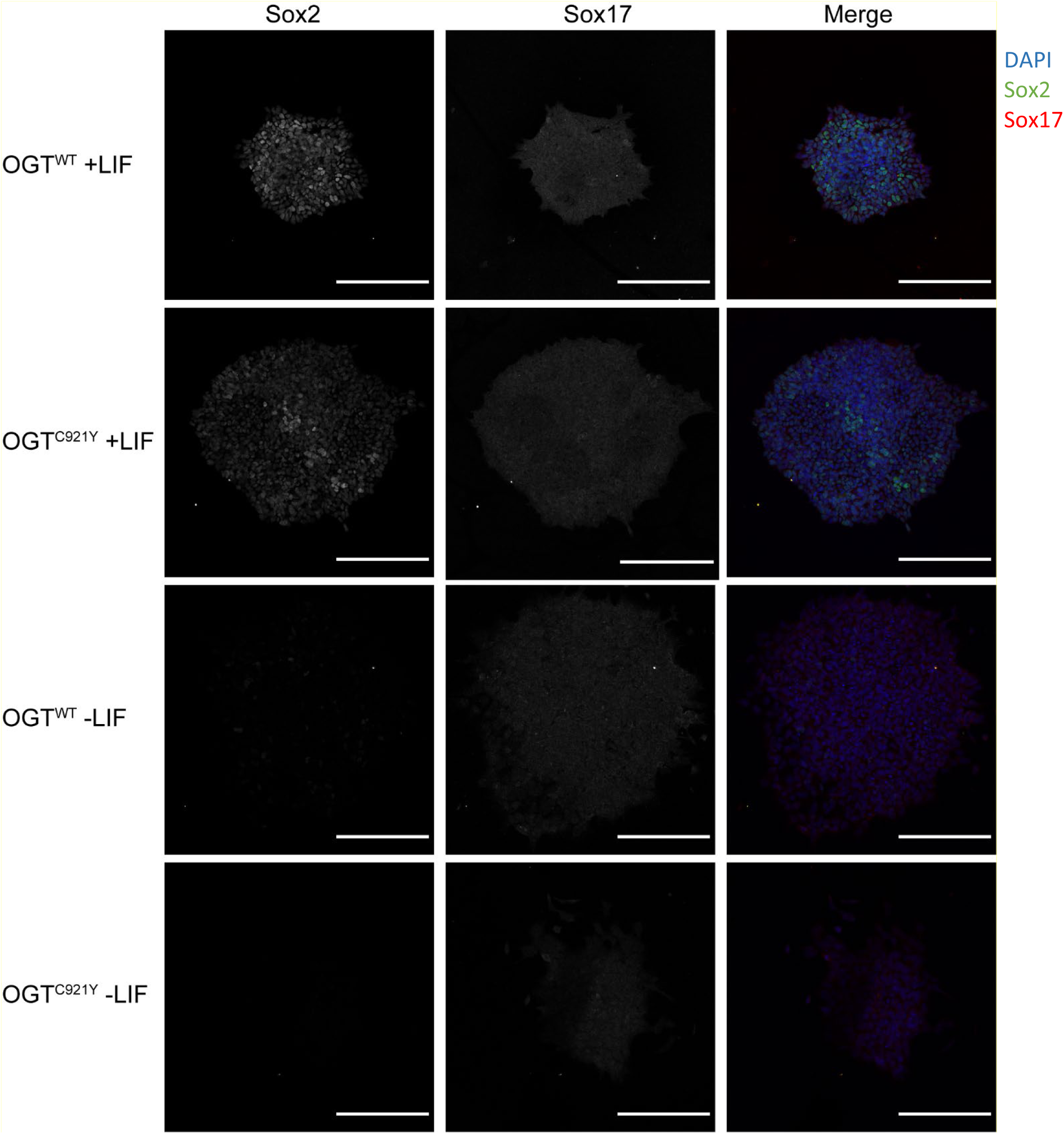
representative images of Sox2 and Sox17 staining in OGT^WT^ and OGT^C921Y^ colonies. Scale bar = 200 µM.

**Supplementary figure S14:**
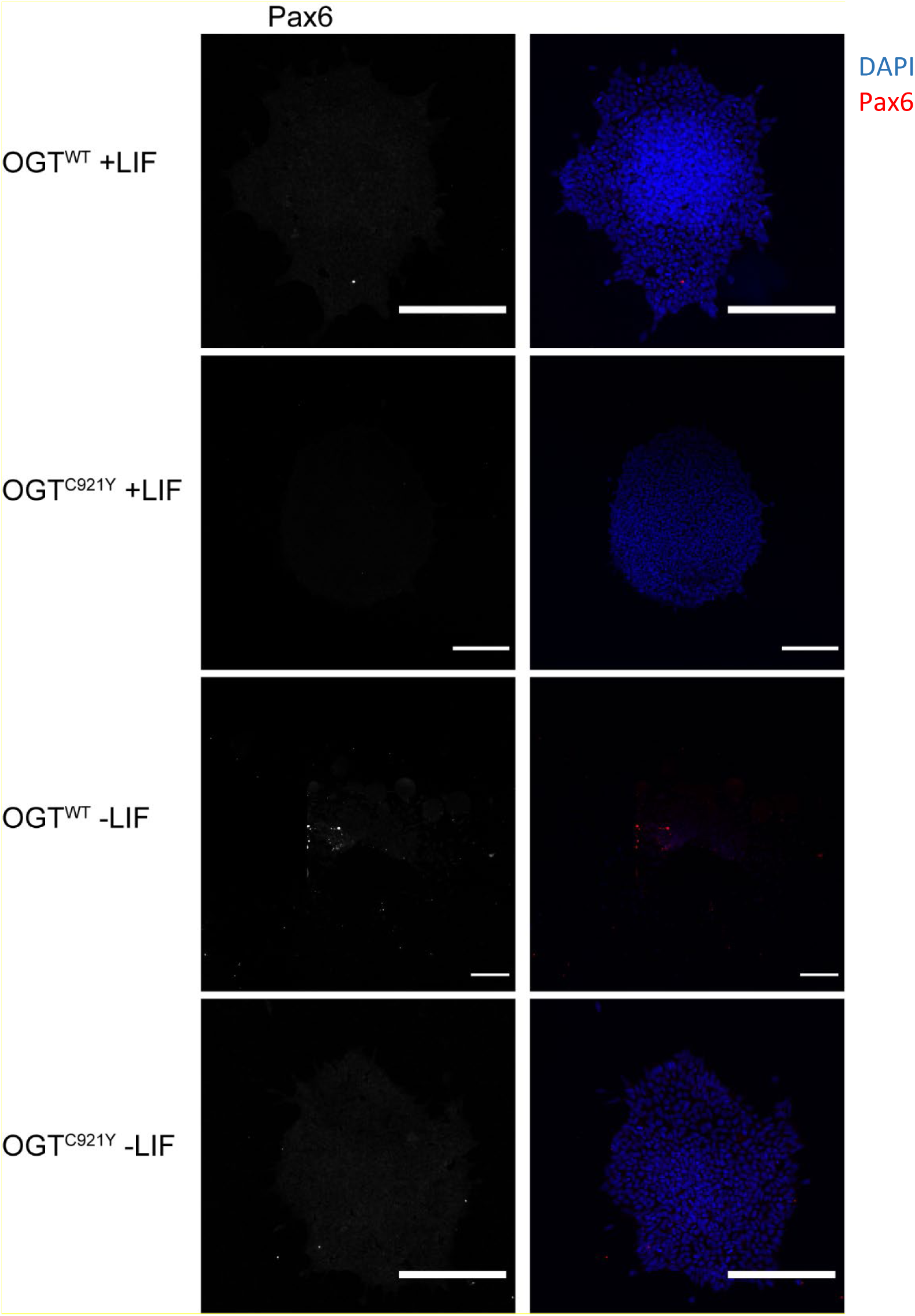
representative images of Pax6 staining in OGT^WT^ and OGT^C921Y^ colonies. Scale bar = 200 µM.

**Supplementary table 1:**
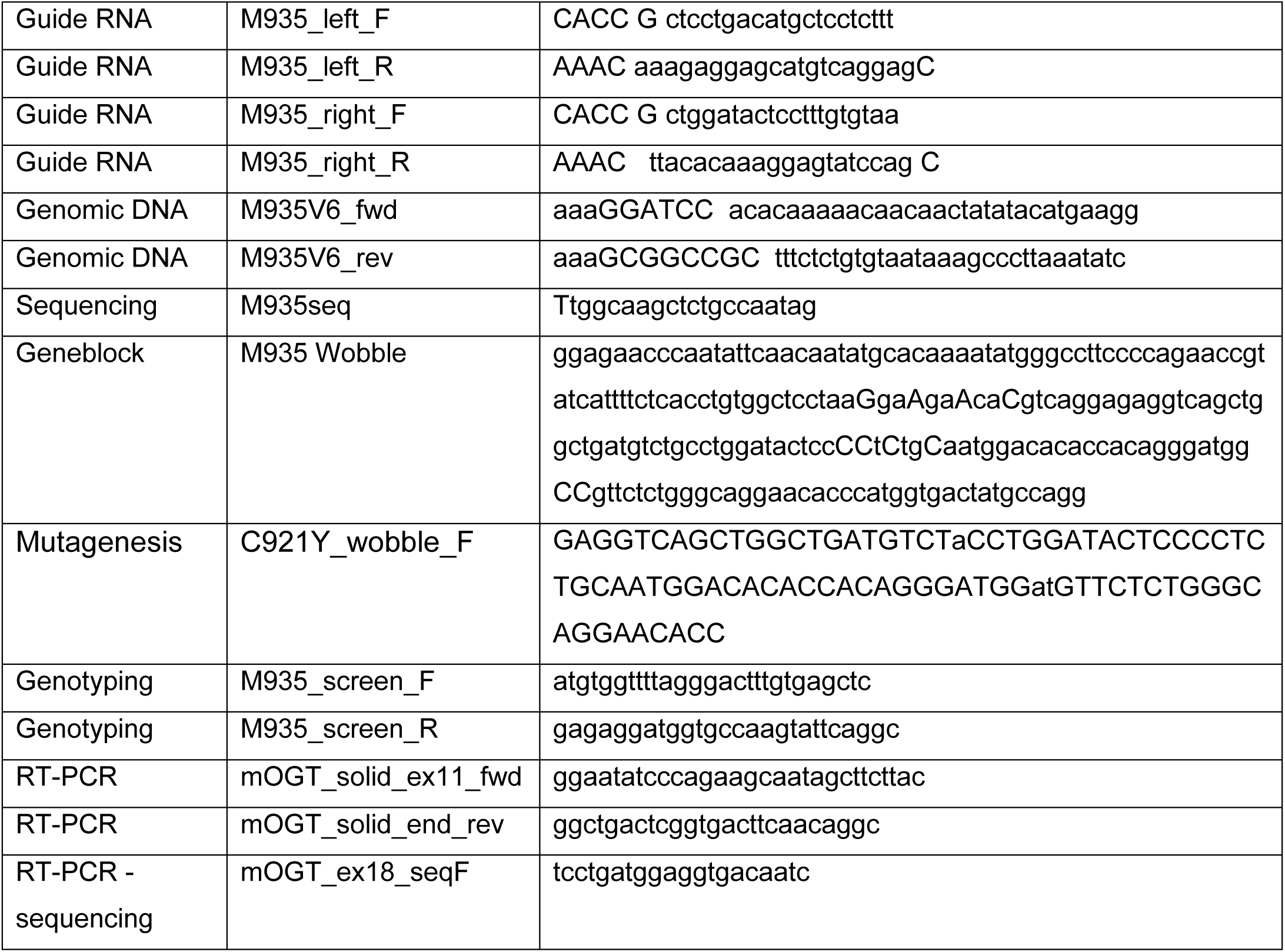
Sequences of custom-made reagents used for generation of OGT^C921Y^ mESC lines.

**Supplementary table 2:**
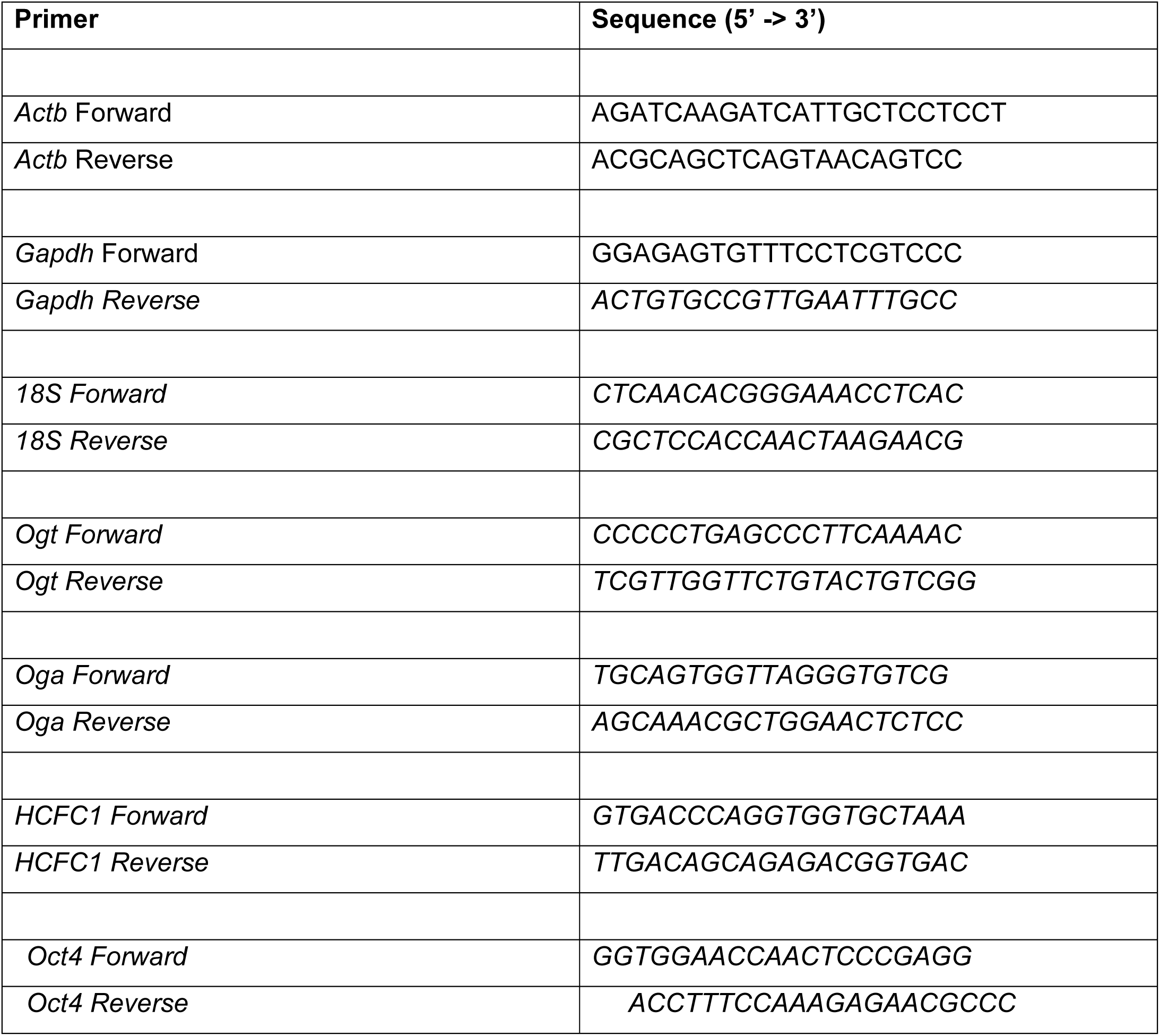
Sequences of qPCR primers.

**Supplementary information 1:**
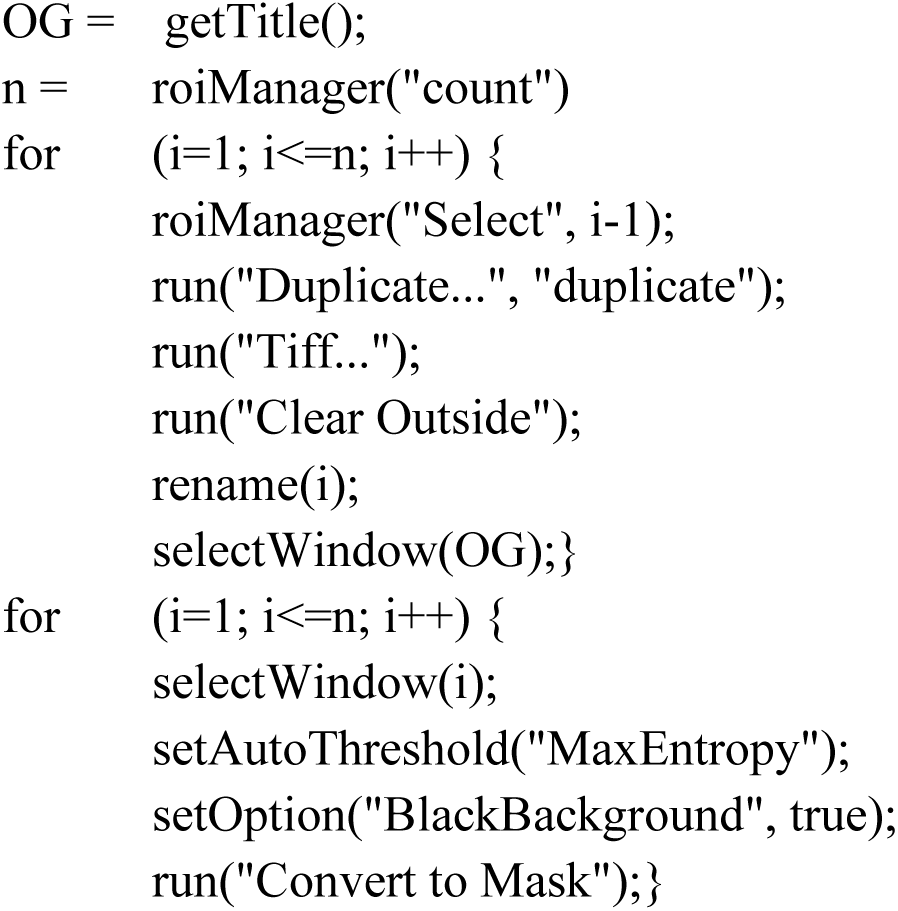
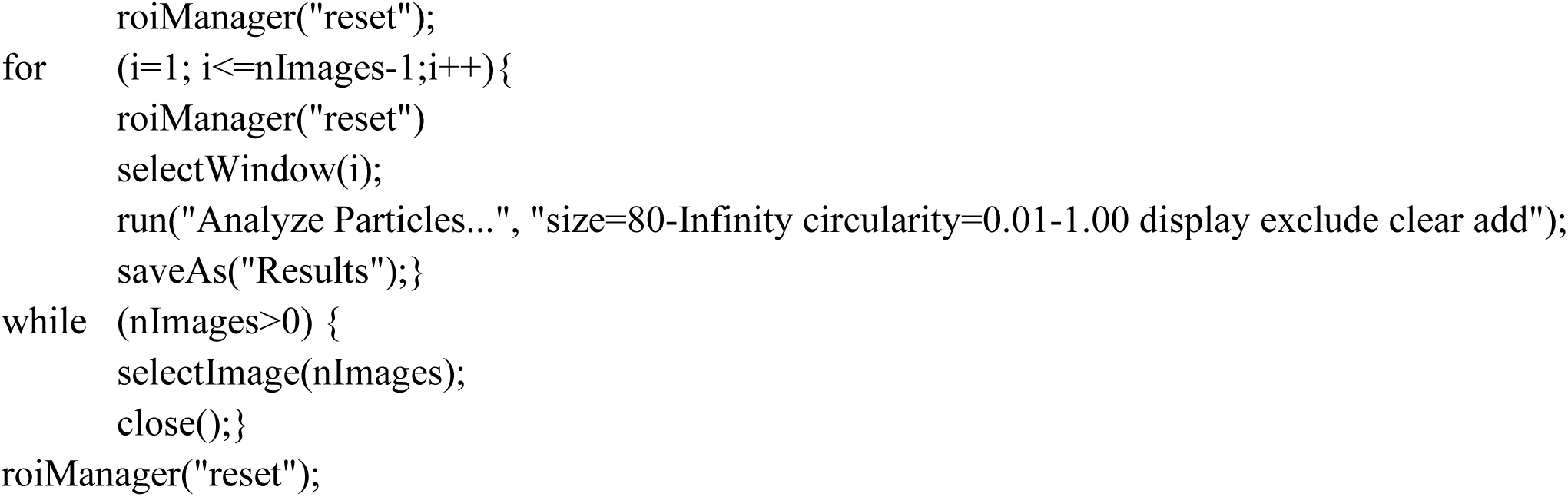
ImageJ colony analysis macro.

**Supplementary information 2:**
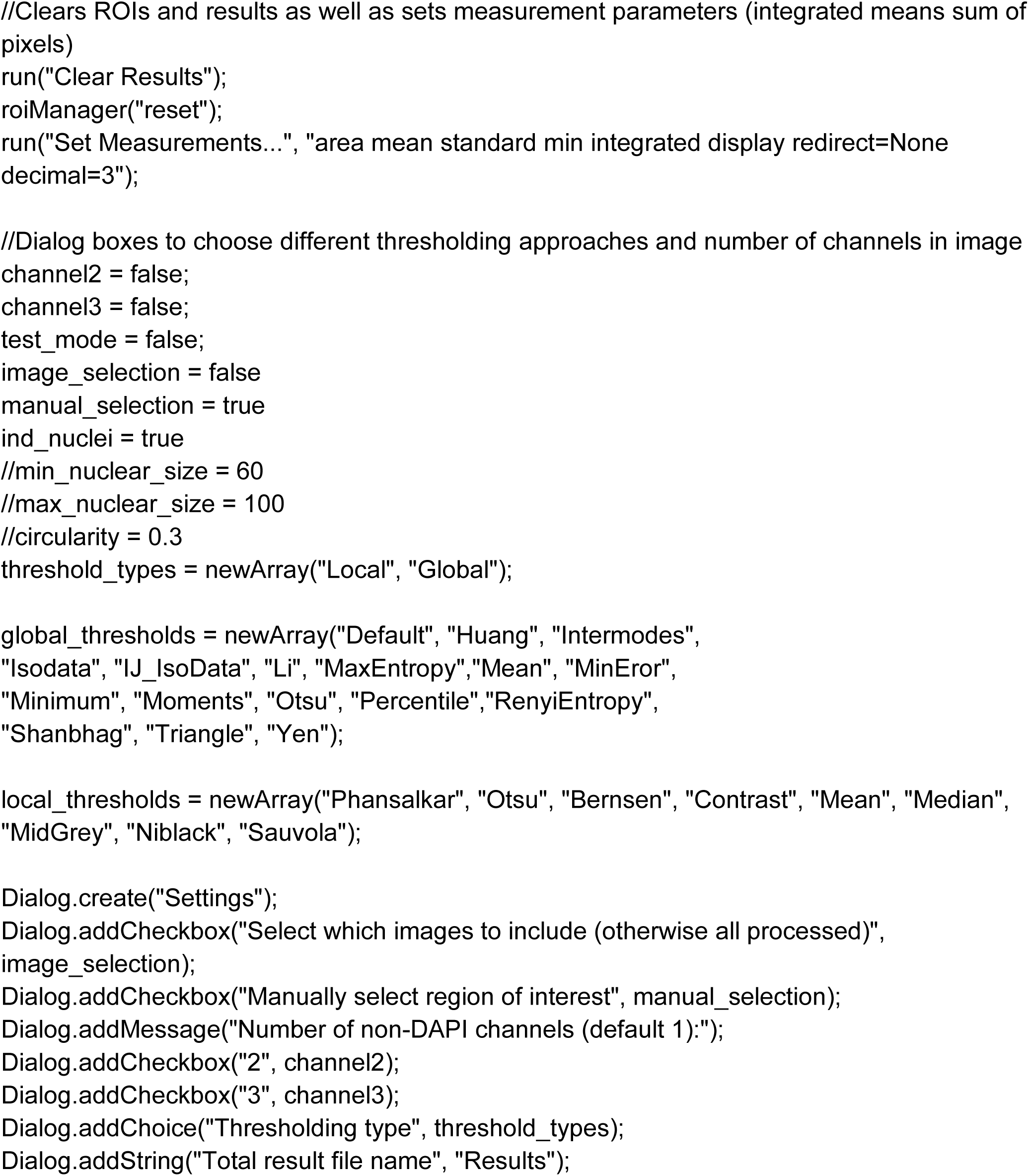
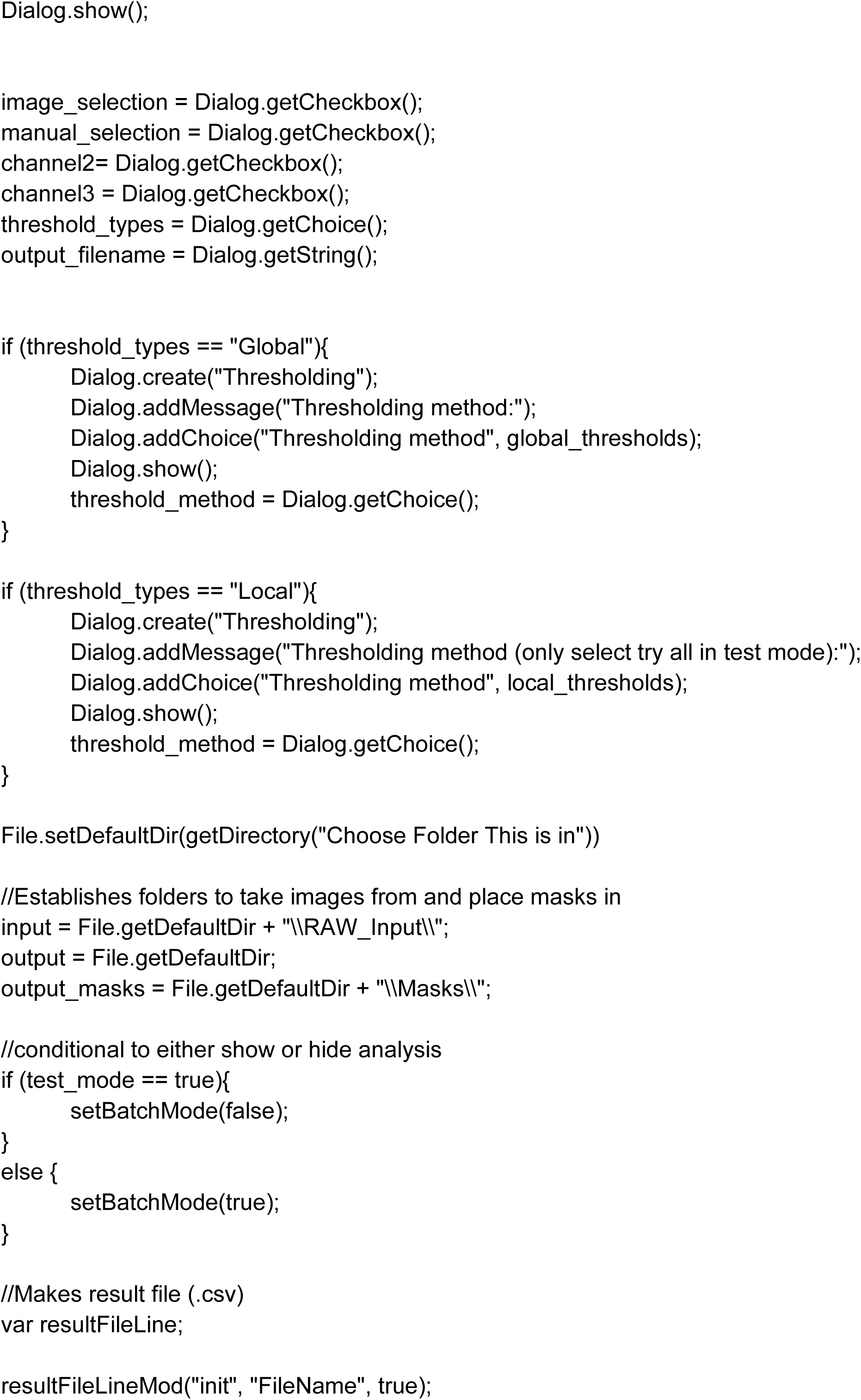
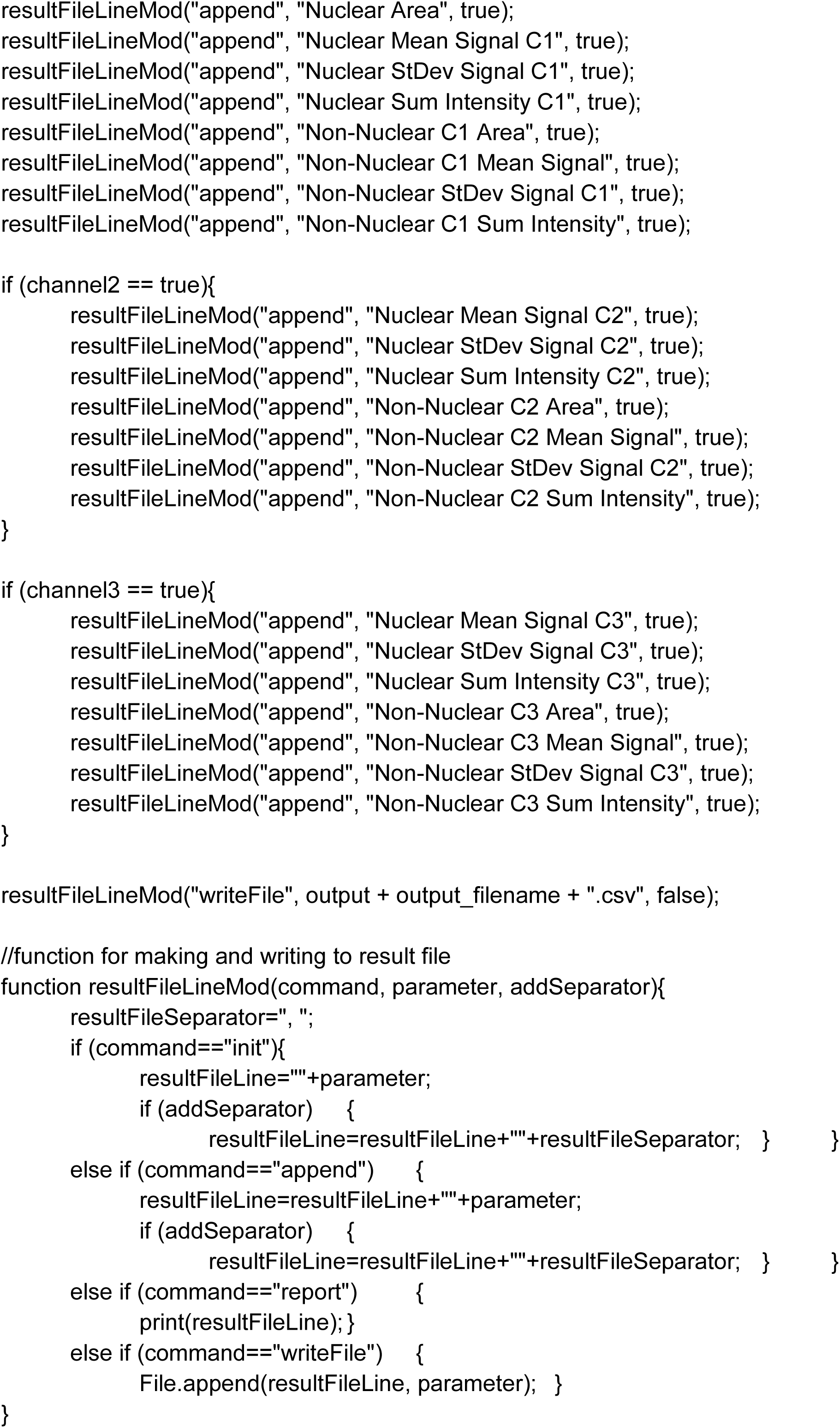
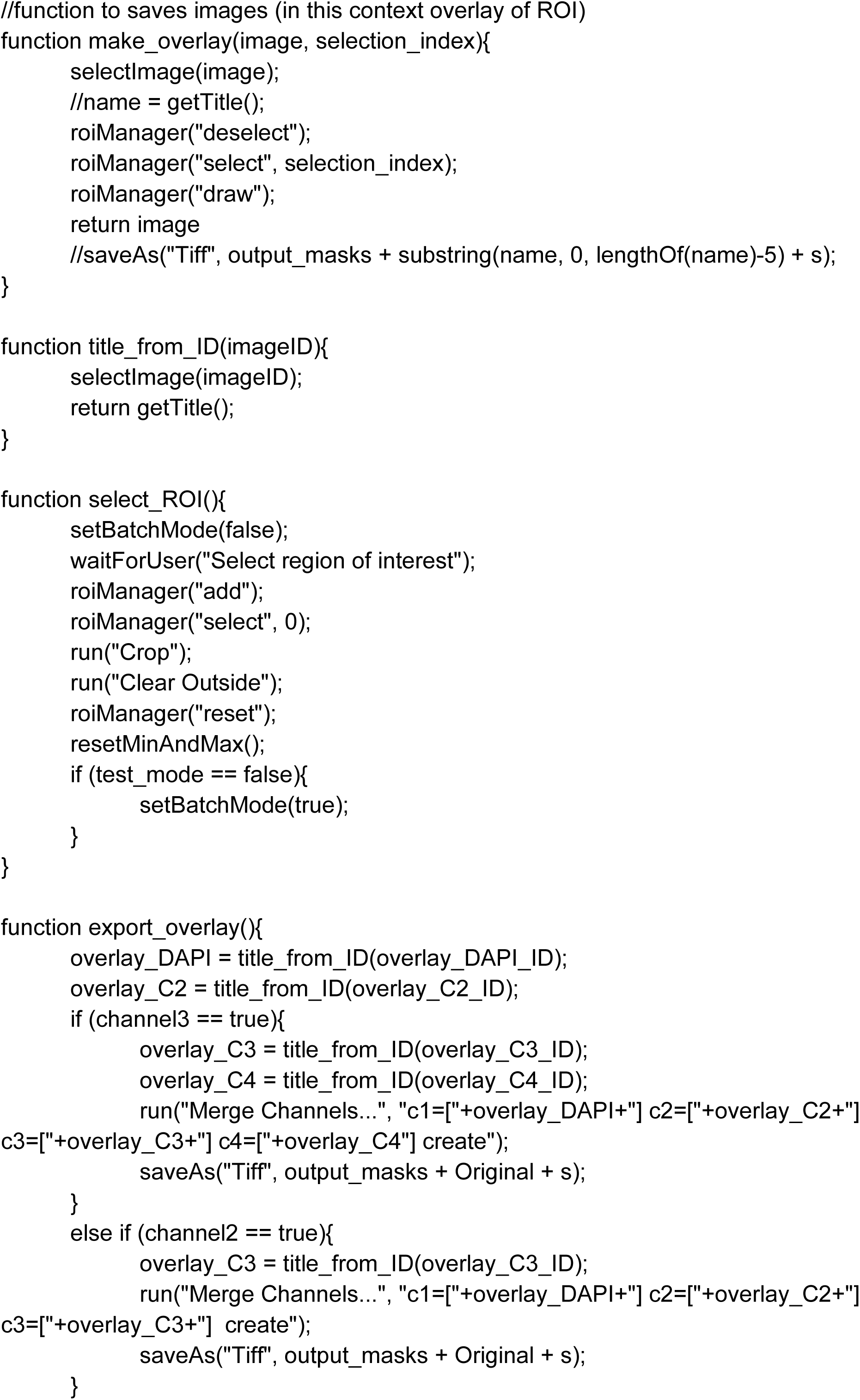
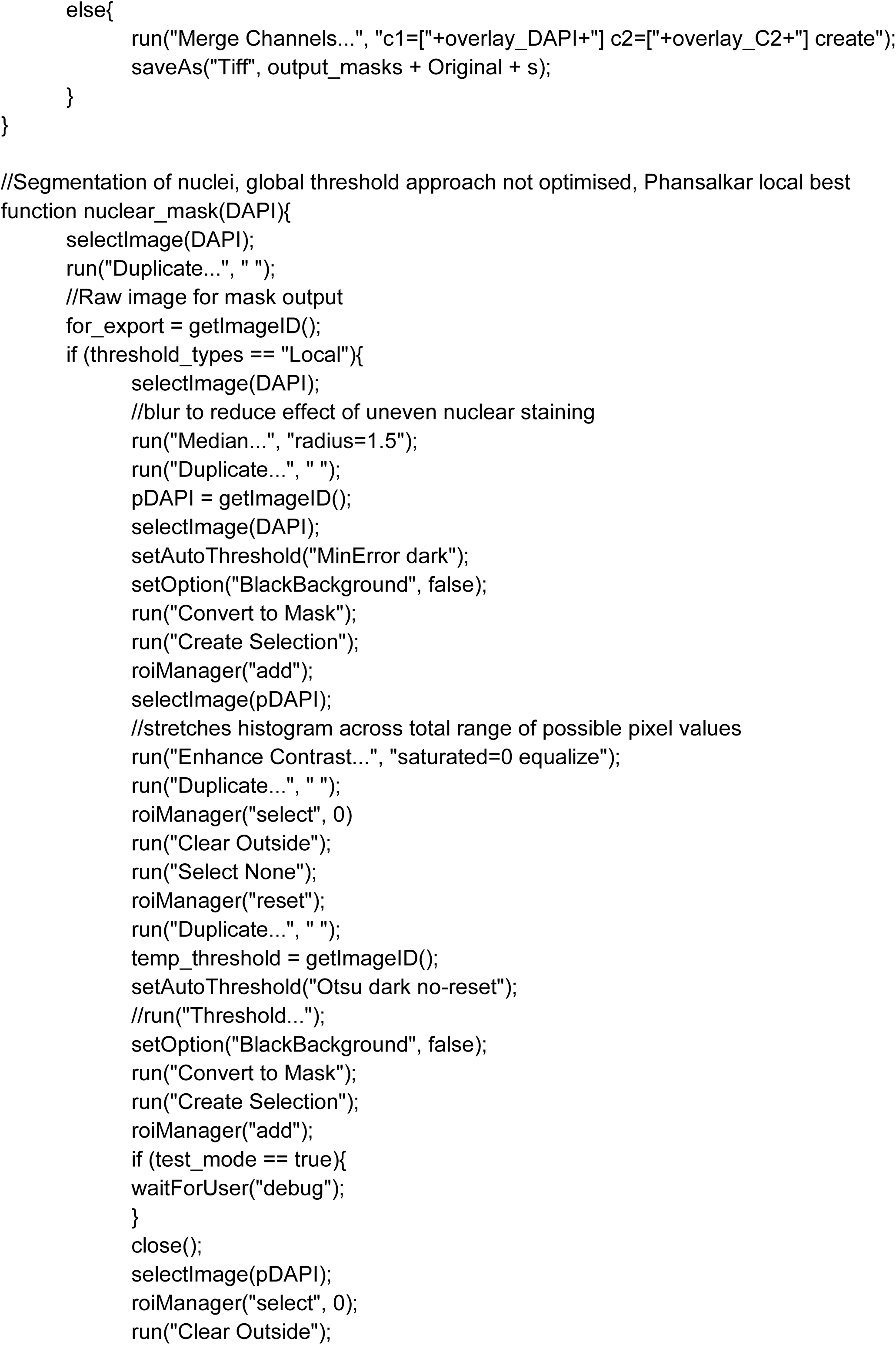
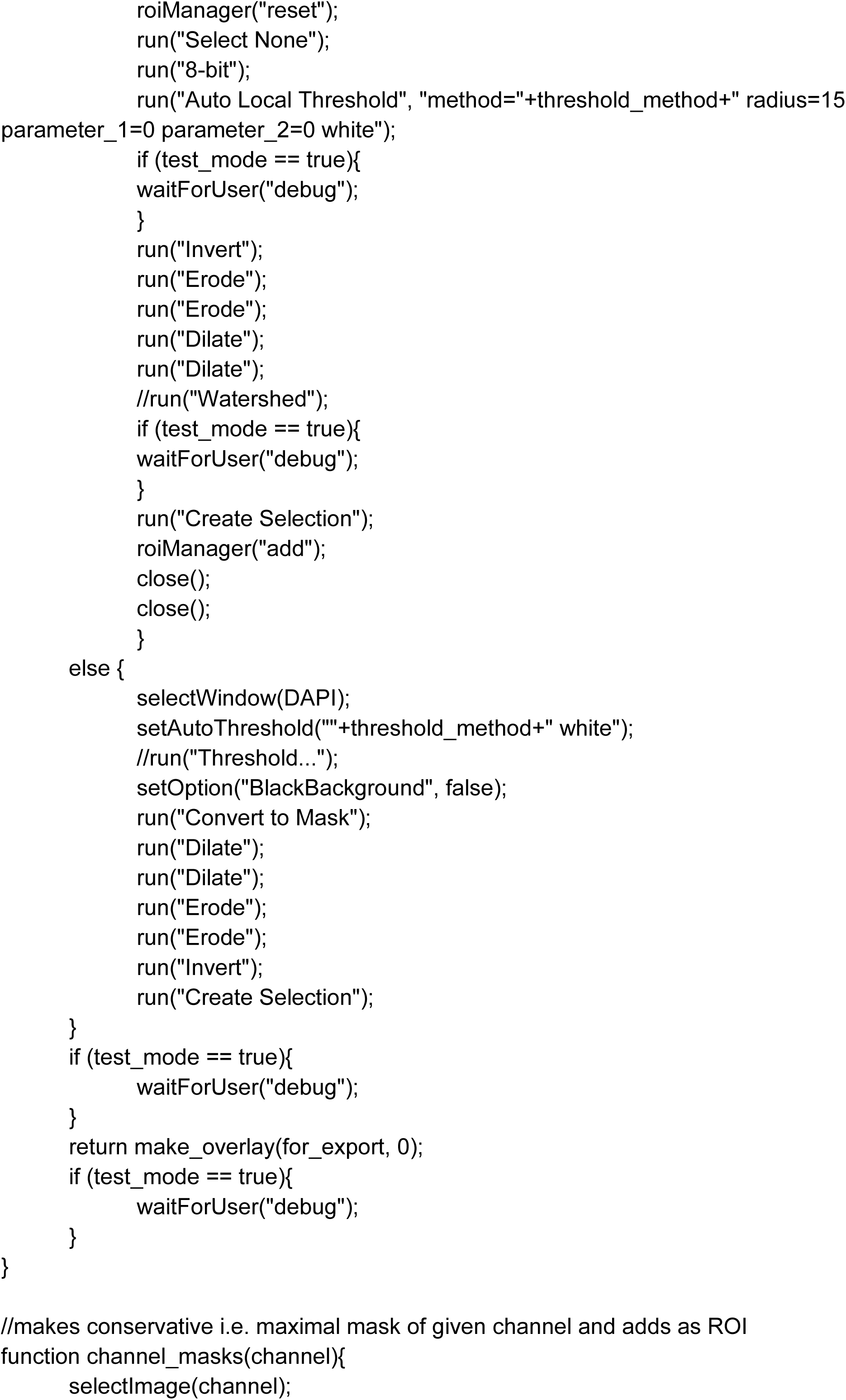
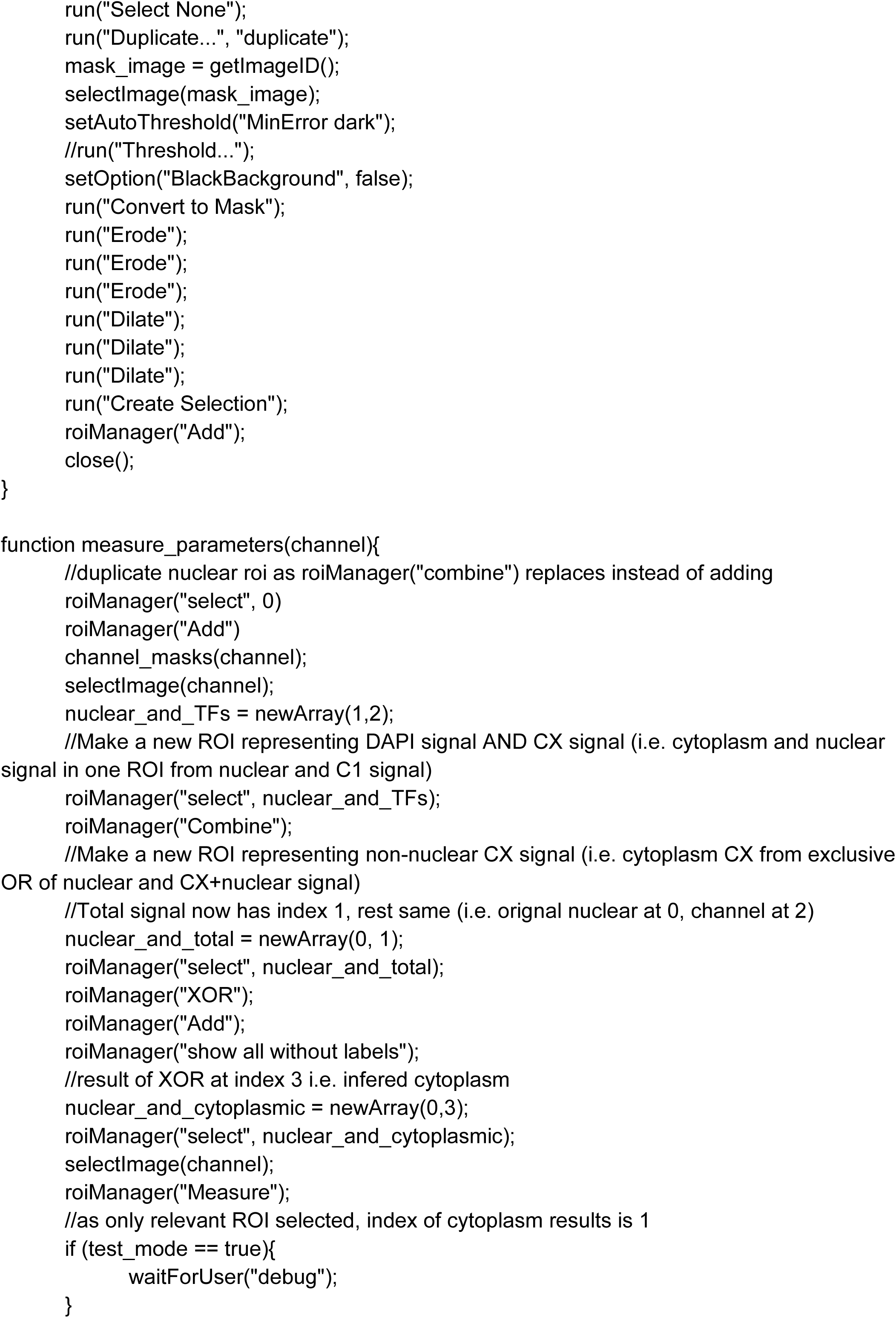
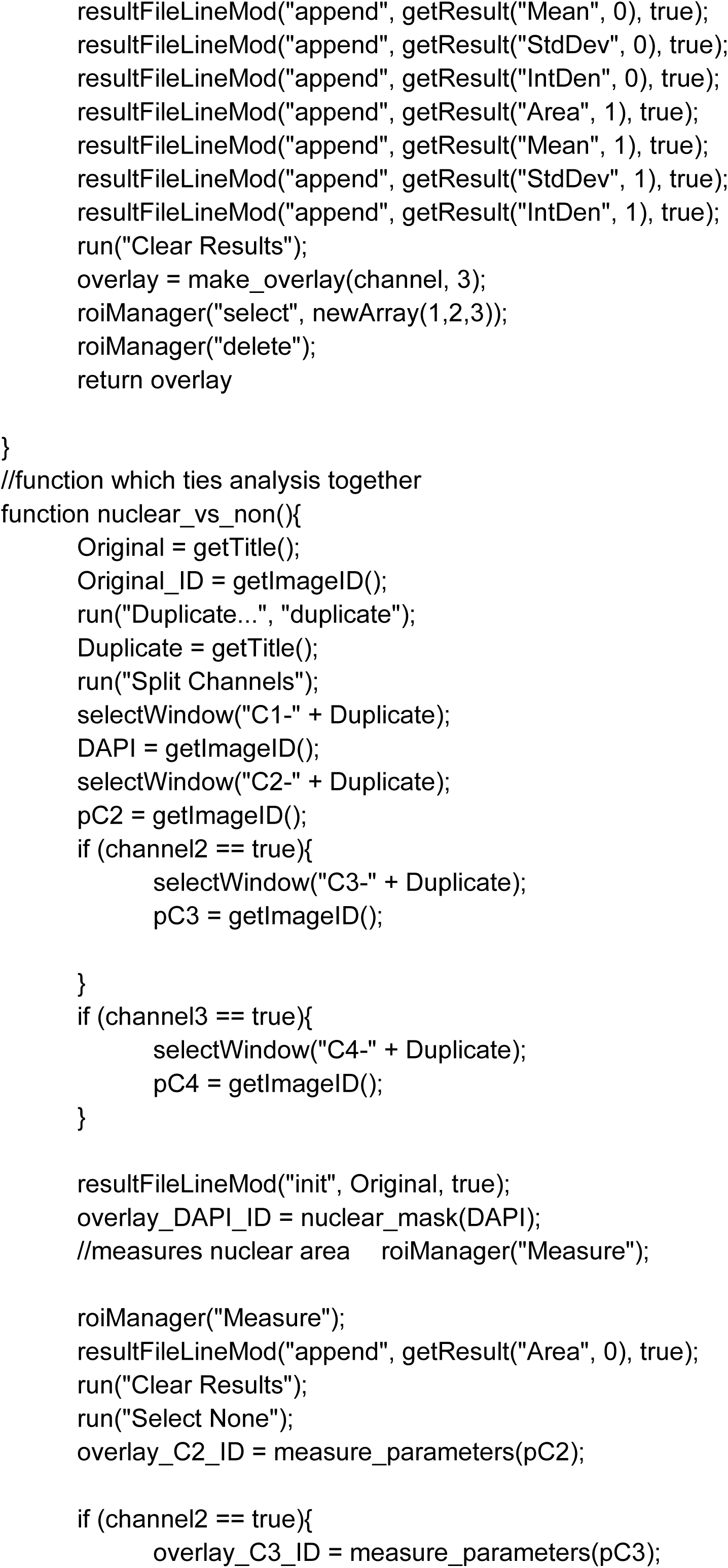
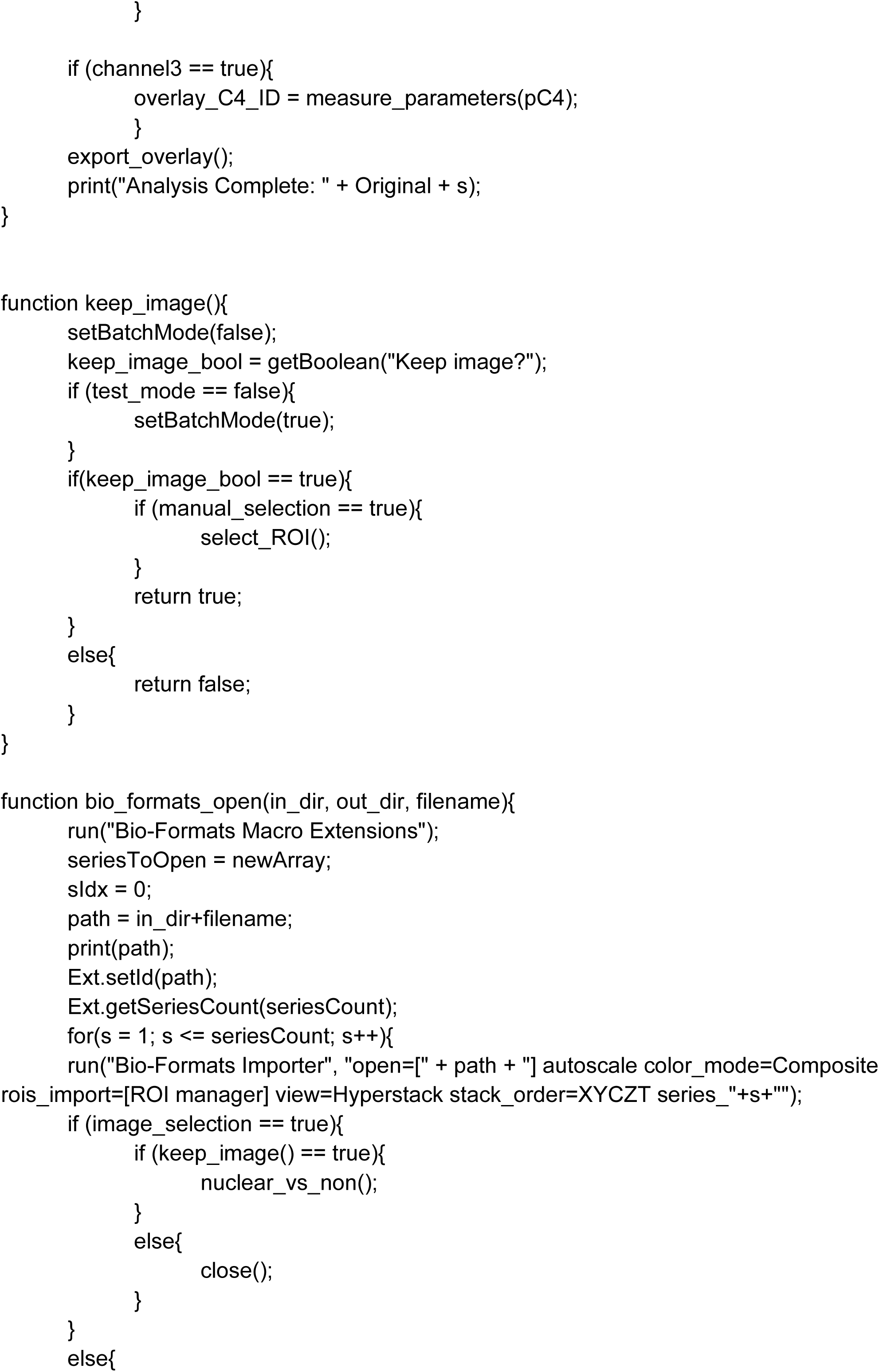
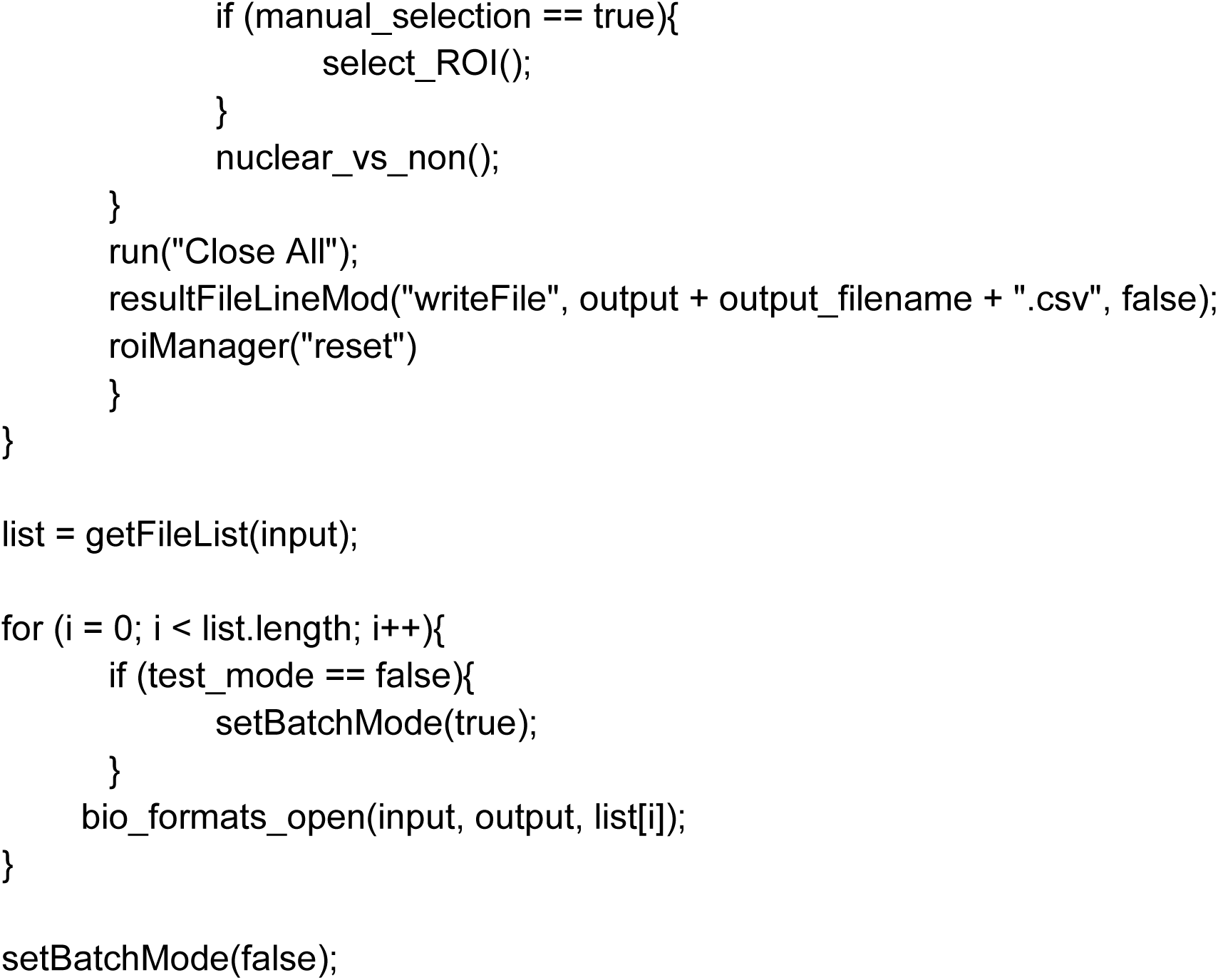
ImageJ colony analysis macro.

